# Source-sink reduction and improvement in rapeseed (*Brassica napus* L.) during the exponential grain filling phase and responses of grain yield, its components and grain quality traits

**DOI:** 10.1101/2025.10.17.683112

**Authors:** Sebastian Garcia, Jose F. Verdejo, Daniel F. Calderini

## Abstract

Rapeseed (*Brassica napus* L.) final grain weight and in turn grain yield, results from the interaction between assimilate supply (source) and sink capacity; however, the extent to which source limitation constrains yield formation during grain filling remains under debate. Understanding how the manipulation of the source–sink ratio (S–S ratio) affects yield and grain traits is critical for elucidating the physiological mechanisms behind yield stability in high-yield environments. This study aimed to evaluate how variations in the S–S ratio during the grain-filling phase influence grain weight and yield, biomass allocation, grain-filling dynamics, and grain quality traits in rapeseed. A field experiment was conducted during two seasons in Valdivia, Chile. One high-yield potential and adapted hybrid (Click CL) was evaluated under three radiation regimes in a randomized complete block design: control, −50% incident radiation (shading), and +50 % incident radiation (reflected radiation pannels, PET). S–S ratio treatments were applied from the beginning of grain filling (BBCH 71) to physiological maturity (BBCH 89) aimed at modify the S-S ratio during the actual grain filling period. The reduced S–S ratio increased thousand-grain weight (TGW), particularly in basal siliques, resulting in yield compensation and demonstrating a strong structural and physiological buffering capacity. Conversely, increasing the S–S ratio enhanced grain number and grain yield, while TGW remained stable. Grain quality traits responded asymmetrically: under reduced S–S, oil concentration slightly declined whereas protein concentration increased. The increased S–S ratio, had no effect on grain oil and protein concentrations, remaining similar to the control. Sieving analyses revealed a shift toward larger grain size classes under reduced S– S, whereas the distribution under increased S–S resembled the control. Overall, these findings indicate that rapeseed maintains yield stability through compensatory adjustments in grain weight and size distribution under contrasting assimilate availabilities. Under high-radiation temperate conditions, rapeseed productivity during grain filling is predominantly governed by sink capacity, highlighting its physiological plasticity and resilience to variations in source–sink balance

**Highlights:**

1. In high-yield conditions without structural changes, grain filling depends on sink capacity.
2. A 50% reduction in radiation increases grain weight and maintains grain yield.
3. A 50% increase in radiation raises grain number and yield via more grains per plant.
4. Source reduction shifts grains to larger sizes; source increase maintains stability.
5. Oil in grain is stable with increased radiation, declines when it is reduced.

## 1. Introduction

World population is projected to reach ∼9.7 billion by 2050, which is a major challenge to ensure global food security (Fischer & Edmeades, 2010; United Nations, 2024). In this context, strategic crops like rapeseed (*Brassica napus* L.), a key source of healthy vegetable oil and high-quality proteins, play a crucial role due to their remarkable content of unsaturated fatty acids and well-balanced amino acid profiles, making it highly valuable for human nutrition (Aider & Barbana, 2011; Wu & Muir, 2008). Currently, rapeseed ranks as the third most consumed vegetable oil globally, following palm and soybean oils, with world production reaching 33.1 million tons in 2023 (UFOP, 2024). Beyond its role in food production, the growing demand for biofuels and animal feed, particularly in aquaculture systems such as those in Chile, has further increased its economic relevance. For instance, rapeseed oil is used as an alternative to fish oil in salmon feed, contributing to the sustainability of the aquaculture sector (Bell et al., 2001; Rondanini et al., 2012; Rondanini et al., 2014). In temperate regions such as southern Chile, rapeseed benefits from favorable agroclimatic conditions, including a high photothermal quotient, moderate temperatures, and appropriate supply of rainfall (Mera et al., 2015; Dörner et al., 2009). These environmental conditions allow for high yields in both winter and spring crop cycles, positioning the region as a strategic area for rapeseed production and industrialization (Mera et al., 2015; Verdejo & Calderini, 2020). However, fully exploiting the yield potential of this crop requires a deep understanding of the physiological mechanisms determining grain yield (GY) and quality, particularly during key reproductive stages. Moreover, the actual and potential constraints to rapeseed production are of acute importance.

Historically, breeding efforts in rapeseed and other grain crops have improved GY by the increase of grain number (GN), due to its high responsiveness to breeding and management, whereas grain weight has shown lower variability and higher heritability (Peltonen-Sainio et al., 2007; Sadras, 2007; Calderini & Slafer, 1998). However, oil crops such as sunflower, heavier grains maintaining grain number have been proposed as an effective strategy for breeding (Pereira et al., 1999; Board & Tan, 1995). The positive impact of heavier grains on yield has been documented in various staple crops, including wheat and soybean (Calderini et al., 2021; Borrás et al., 2004), and also in oil crops such as sunflower and rapeseed, where increased grain weight contributes positively to both grain and oil yield (Chay & Thurling., 1989; Tzen et al. 1993 Castillo et al., 2017; Labra et al., 2017; Verdejo & Calderini, 2020).

Final grain weight is the result of two main key processes, (i) the setting of grain weight potential and (ii) the filling of growing grains (Fig. 4 in Calderini et al., 2001; Egli, 2004). On these regards, it has been demonstrated that the setting of grain weight potential occurs before grain growth, which is supported by the association between final grain weight and the ovary size at pollination across several crop species (Schou et al., 1978; Scott et al., 1983; Calderini et al., 2001; Hasan et al., 2011; Lizana & Calderini, 2013; Castillo et al., 2017). Nonetheless, different physiological regulations of grain filling have been reported among crops. In wheat (*Triticum aestivum* L.), there is general agreement that grain growth is not limited by current photosynthesis, due to the strong buffering capacity by stem reserves (Slafer & Savin, 1994; Gan et al., 2008; Borrás et al., 2004; Savin et al., under review). In contrast, in maize (*Zea mays*) source limitation has been highlighted when assimilates availability is reduced during the linear grain-filling phase (Borrás et al., 2004; Abeledo et al., 2020). On the other hand, the response to the source-sink (S-S) balance in rapeseed during grain filling remains controversial, where some studies have indicated that grain weight is constrained by source limitation (e.g., Fortescue & Turner, 2007; Iglesias & Miralles, 2014), while other experimental results suggest that the sink capacity, defined by ovary size and silique architecture, is the primary determinant of final grain weight (e.g., Tayo & Morgan, 1975; Tayo & Morgan, 1979; Wang et al., 2011; Li et al., 2018).

Across crop species, the study of grain filling can be approached through the concept of the S-S balance, formalized by Slafer and Savin (1994), which builds upon earlier studies. Rawson and Evans (1971) analyzed the mobilization of stem reserves in wheat, highlighting the role of pre-anthesis assimilates in supporting grain growth. Subsequently, Evans (1984) evaluated the effects of irradiance on GY and its components by reducing incident solar radiation by approximately 50% during the critical period for grain number determination (see Fischer, 1985; Kantolic & Slafer 2004). These studies set the foundations for understanding how the interaction between the source capacity and the sink strength determines final grain weight. This physiological relationship is particularly relevant in rapeseed, where yield components are defined during flowering and grain-filling stages (Keiller & Morgan, 1988; Diepenbrock, 2000; Zhang & Flottmann., 2018).

The potential number of siliques is set prior to flowering through the formation of floral primordia, while their final number depends on assimilate availability both before and during flowering (Habekotté, 1993; Poggio et al., 2004). Through these phenological phases, GN and potential grain weight are setting, especially during the ‘critical period’ for GN, which spans from 100 to 500 °Cd after the starting flowering (Kirkegaard et al., 2018). However, the effect of S–S manipulations during the grain filling period (BBCH 70–89) remains controversial in rapeseed. Some modeling studies suggest a progressive decline in the S-S ratio due to the high energetic costs of maintaining a complex reproductive architecture, which may reduce the biomass partitioning efficiency (Jullien et al., 2011; Verdejo & Calderini, 2025). Experimental results on S–S manipulations in rapeseed were often inconsistent, partly due to the diversity of methodologies employed. These manipulations can be classified into structural (vegetative or reproductive) and environmental categories, each affecting GN, thousand grain weight (TGW) and GY in different ways (Table A1). To facilitate the interpretation, the reported results were grouped according to the type of manipulation and whether the effect was negative, neutral or positive. When the source was reduced through defoliation, shading or by increased plant density, grain weight consistently decreased, with TGW reductions from 9 to 18% and up to 40% in GN. For instance, basal defoliation (BBCH 71–75) reduced TGW by 12% and GN by 18%, while increased plant density (50– 60 plants m⁻²) lowered TGW by 18% without affecting GN. Under continuous shading from flowering to maturity (BBCH 60–89), total yield declined up to 15%, with TGW decreased by 12% and oil content also negatively affected. These results indicate that source limitation during grain filling tends to reduce grain size more than grain number, particularly when the S-S manipulation was applied during BBCH 70–89.

In contrast, when the S–S ratio was increased, either by partial silique removal or by removing late-developing branches, compensatory effects were often observed. For instance, TGW increased by 10– 15%, and in some cases GN per plant also rose up to 30% (Brar & Thies, 1977; Williams & Free, 1979; Yates & Steven, 1987; Rao, 1991; Diepenbrock, 2000; Kirkegaard et al., 2012). Notably, the removal of basal siliques early during grain filling (BBCH 69–75) led to an extended flowering duration and the development of additional fertile flowers, partially offsetting sink reduction (Peltonen-Sainio et al., 2007; Takashima et al., 2013; Zhang & Flottmann., 2018). These outcomes suggest that reproductive plasticity can buffer yield loss under mild sink limitation (Table A1).

Neutral responses were also reported. In several cases, manipulations such as light competition or partial defoliation under water stress reduced TGW by ∼15% but did not significantly affected GN. Some genotypes with open canopies (e.g., apetalous lines) maintained high GN and TGW despite reduced leaf area, likely due to improved light penetration and assimilate distribution (Table A1). In addition, to have a precise evaluation of the source-sink balance during grain filling in oil crops, the quantification of the impact of source-sink manipulation on grain oil, besides grain weight, is a key for this analysis.

Overall, most studies evaluated the S–S ratio through indirect structural or environmental alterations, such as defoliation or shading, rather than by directly manipulating the amount of intercepted radiation. In contrast, the present study isolates this factor by modulating solar radiation interception during the linear phase of grain filling (BBCH 71–89), allowing a more precise evaluation of how decreased or increased solar radiation affects GY, GN, TGW, canopy biomass partitioning, and grain composition under the high-yield condition of southern Chile. Recently, Savin et al. (under review) have critically analyzed the results of source-sink experiments concluding that “*source limitation is not a general condition for effective grain filling, although exceptions may occur*”. We take this reasoning and analysis in mind to assess our objective of evaluating the impact of decreased and increased S–S manipulation on GY, yield components, and quality traits (grain oil and protein concentrations) of rapeseed.

## 2. Materials and methods

### 2.1 Field experiments, experimental design and set up

A field experiment was conducted over two consecutive seasons (2022-23 and 2023-24) at the Experimental Station of the Universidad Austral de Chile, located in Valdivia, Chile (39°47′ S, 73°14′ W), in a Duric Hapludand soil. The high yielding rapeseed hybrid Click CL (Deutsche Saatveredelung AG, Germany) was evaluated under three source-sink ratio (S-S) treatments: (i) control, (ii) decreased S-S, and (iii) increased S-S.

The experiment was arranged in a randomized complete block design (RCBD) with four replicates in each season. Each plot measured 1.5 m in length and 2.5 m in width, with a planting density of 45 plants per m², distributed across 11 rows spaced 0.2 m apart. Sowing dates were September 3 in 2022 (Season 1) and September 28 in 2023 (Season 2).

Previous to sowing, soil samples and the corresponding soil analysis were carried out to evaluate the fertilization requirements. In both seasons, agricultural lime was applied as a preventive measure to counteract soil acidity, as described by Labra et al. (2017) and Verdejo & Calderini, 2020. Nitrogen fertilization was split into two applications of 100 kg N ha⁻¹ each, i.e. the first was applied after plant emergence (BBCH 10), and the second when the fifth internode expanded (BBCH 35), in both seasons. Additionally, 150 kg ha⁻¹ of Ca(H₂PO₄) ₂ was applied at sowing to prevent nutritional phosphorus deficiencies. Sulfur (S), magnesium (Mg), and potassium (K) were also supplied at sowing using Sulpomag® as fertilizer.

To avoid biotic constraints from pests, diseases, and weeds; optimal management practices were implemented following manufacturer recommendations. Specific practices included Gaucho® 600 FS insecticide-treated grains for early pest protection and Basamid® herbicide application before sowing to control weeds. Plots were irrigated using a drip irrigation system throughout the crop cycle in both seasons to supplement natural rainfall, ensuring adequate water availability until physiological maturity (BBCH 87).

### 2.3 Source-sink treatments

S–S ratio treatments were applied from the beginning of grain filling (BBCH 71) until physiological maturity (BBCH 89), in order to avoid interference with the critical period of grain number determination (BBCH 60–69, Kirkegaard et al., 2018). These treatments were specifically designed to be effective during the linear phase of grain filling, when the assimilate accumulation in developing grains occurs at its highest rate. The control treatment was conducted without any S-S manipulation throughout the crop cycle. The reduced S–S ratio treatment consisted of a wooden-frame structure covered with black Rachel mesh, which reduced incident solar radiation by 50%. The mesh was positioned 0.3–0.4 meters above the canopy, covering 6.5 m^2^ per plot, and its south side was left open to facilitate air circulation and pollinator access, following previous field protocols at EEAA (Labra et al., 2017; Verdejo & Calderini, 2020). In contrast, the increased S–S ratio treatment involved six reflective panels placed along the south side of each plot, oriented east to west and inclined at 45° and 60°, designed to increase radiation availability by approximately 50%. Each panel was built using a wooden frame and a reflective surface made of cardboard coated with polyethylene terephthalate film, a material with high optical clarity, low absorption coefficient, and a refractive index of 1.63 in the visible spectrum (Elman et al., 1998). The panels were installed at a height of 1.0 meters above ground level and anchored to vertical posts to ensure structural stability. Figures 1A and 1B show the spatial arrangement and functional layout of the solar radiation reduction and increase structures, respectively, while Figures A2 and B2 depict their three-dimensional configurations at 1:50 scale, illustrating their geometry, proportionality, and their integration within the experimental field. Both structures were designed with strict dimensional accuracy and documented through scaled planimetric and volumetric models to ensure reproducibility and consistency under field conditions.

**Figure 1.**
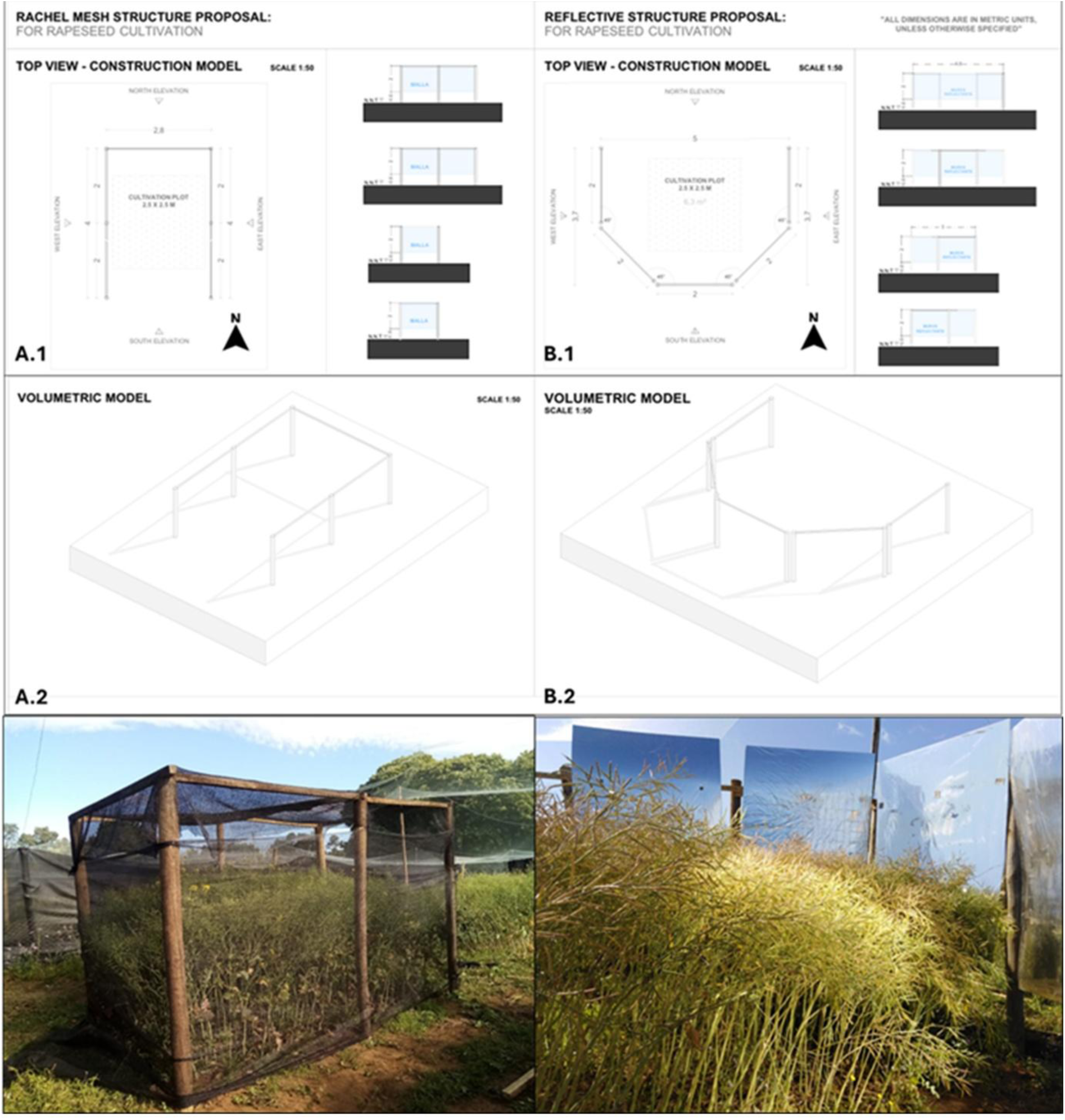
Schematic representation of source–sink ratio (S–Sratio) treatments. Panels A1 and A2 correspond to the source reduction treatment: A1 shows the schematic layout using shading structures (black mesh), and A2 shows its three-dimensional implementation in the field. Panels B1 and B2 correspond to the source increase treatment: B1 illustrates the schematic design with reflective panels, and B2 depicts the volumetric structure used under field conditions. All structures are shown at scale (1:50) to represent their morphology, spatial arrangement, and proportional integration with the crop canopy.

### 2.3 Weather measurements

Meteorological data of temperature, along with incident solar radiation, were measured every 30 minutes from sowing to harvest at the Austral meteorological station of EEAA (http://agromet.inia.cl/), located 250 m from the experiment. The photothermal quotient was calculated by dividing daily solar radiation by the mean daily temperature, with a base of 0°C, a value recommended for rapeseed studies (Adamsen and Coffelt, 2005; Kirkegaard *et al*., 2012) (Fig. 2). Canopy temperature and Photosynthetically Active Radiation (PAR) reaching the crop were measured at each plot. Canopy temperature was measured using two methods: a FLIR E8 Pro thermal imaging camera (FLIR Systems, Inc., Wilsonville, OR USA) focusing on the siliques and branches three times a day (10:00, 13:00 and 15:00 hours), and a digital thermometer positioned at the height of branch 1, recording temperature continuously from 8:00 AM to 8:00 PM. The thermal images were analyzed using FLIR Tools software, where eight points corresponding to siliques and branches were selected in each image to generate high-resolution heat maps (Fig. 3). To measure PAR in the experiment, a LI-191 R sensor (LI-COR Inc.) was positioned 10 cm above the canopy for incident radiation (PAR_inc_) and at different heights within the canopy (at 0.50; 0.75 and 1.0 m above ground level) for transmitted radiation (PAR_trans_), following the methodology described by Labra et al. (2017). Measurements were taken at six points per plot three times a day (10:00, 12:00 and 16:00 hours) on sunny days. These data were analyzed using R software, specifically the ggplot2 and viridis packages, to generate detailed heat maps of PAR distribution The maps incorporated measurements at ground level and at 0.50, 0.75, and 1.00 m, providing a comprehensive visualization of spatial radiation patterns within the canopy (Fig. 4).

**Figure 2.**
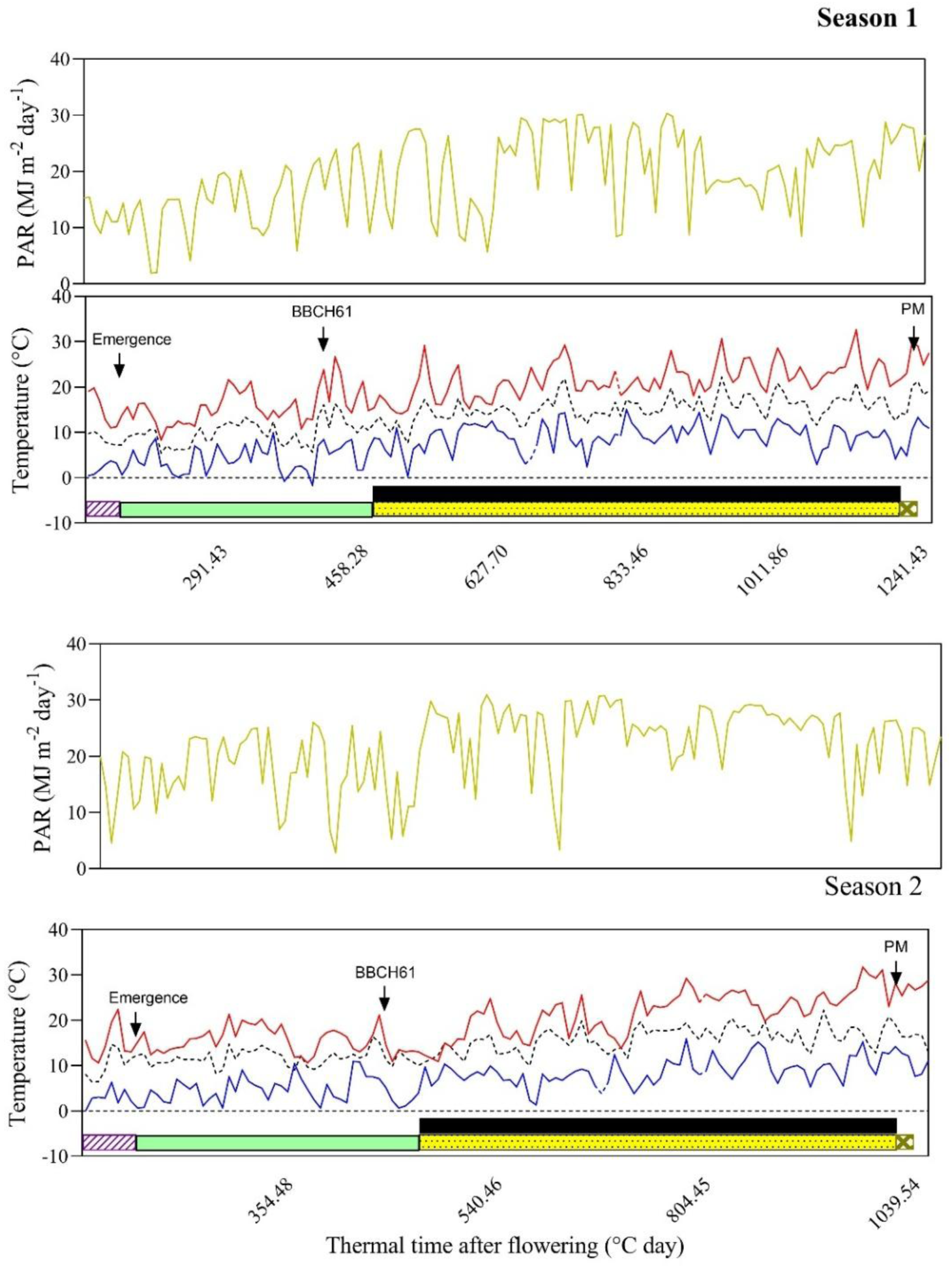
Photosynthetically active radiation (PAR, yellow line), mean daily temperature (black dotted line), maximum daily temperature (red line), and minimum daily temperature (blue line) across the phenological stages. The horizontal bars represent crop phenology from sowing to harvest, divided into the following phases: sowing to emergence (striped, purple bars, BBCH 00–09), emergence to flowering (solid green bars, BBCH 10–61), flowering to physiological maturity (yellow dotted bars, BBCH 61– 87), and physiological maturity to harvest (brown mesh bars, BBCH 87–99) of control treatment in the experiments 1 and 2. PM indicates physiological maturity. Black bars show the period of source–sink ratio (S–Sratio) treatments (BBCH 71–89).

**Figure 3.**
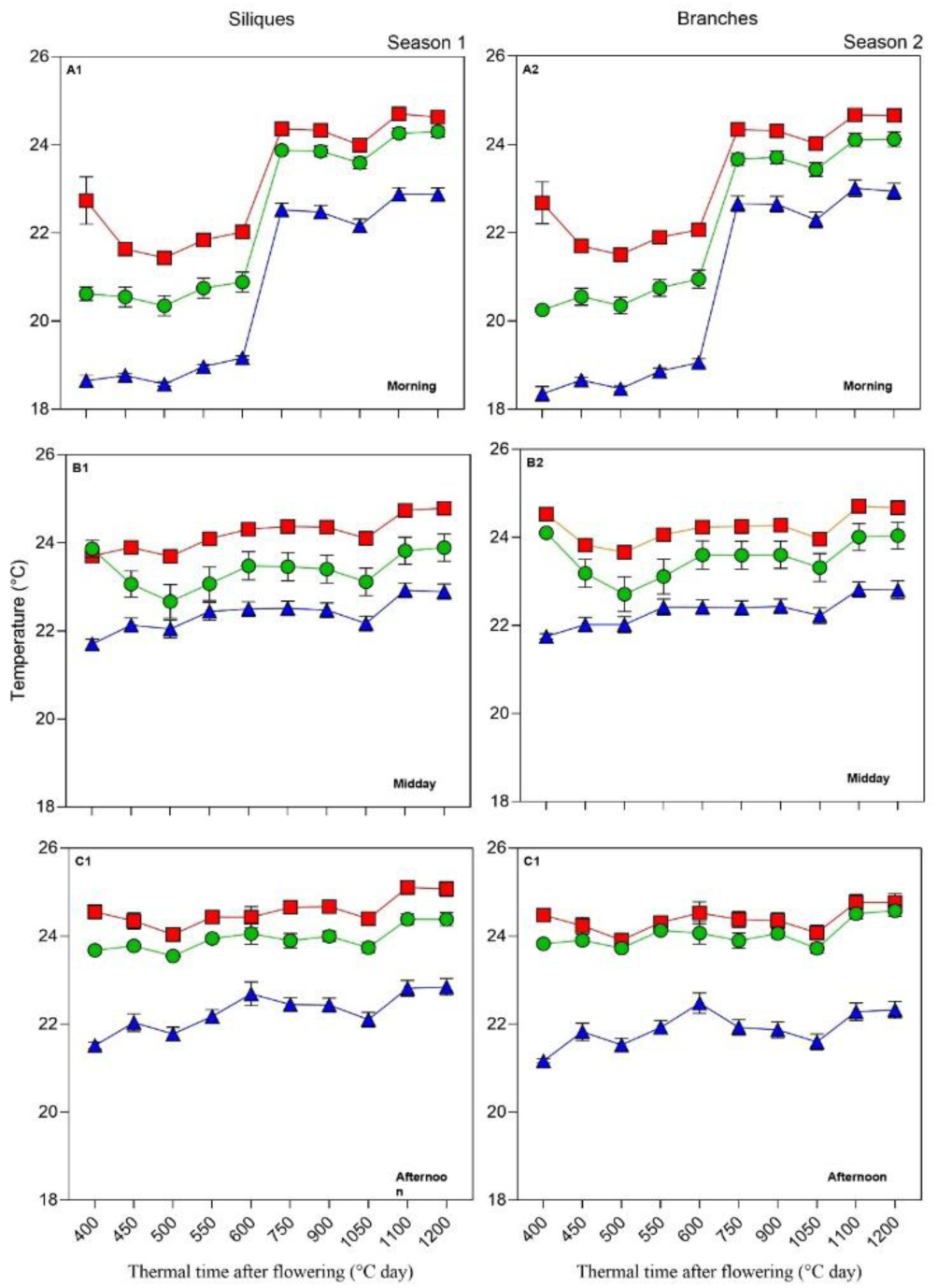
Daily temperatures at three times of the day (morning: 10:00 h, midday: 13:00 h, and afternoon: 15:00 h) measured on siliques (A1, B1, C1) and branches (A2, B2, C2) from the end of flowering (BBCH 69) to harvest (BBCH 89) during the experiment. Treatments are represented by colors and symbols: control (green triangles), source increase (red squares), and source decrease (blue circles).

**Figure 4.**
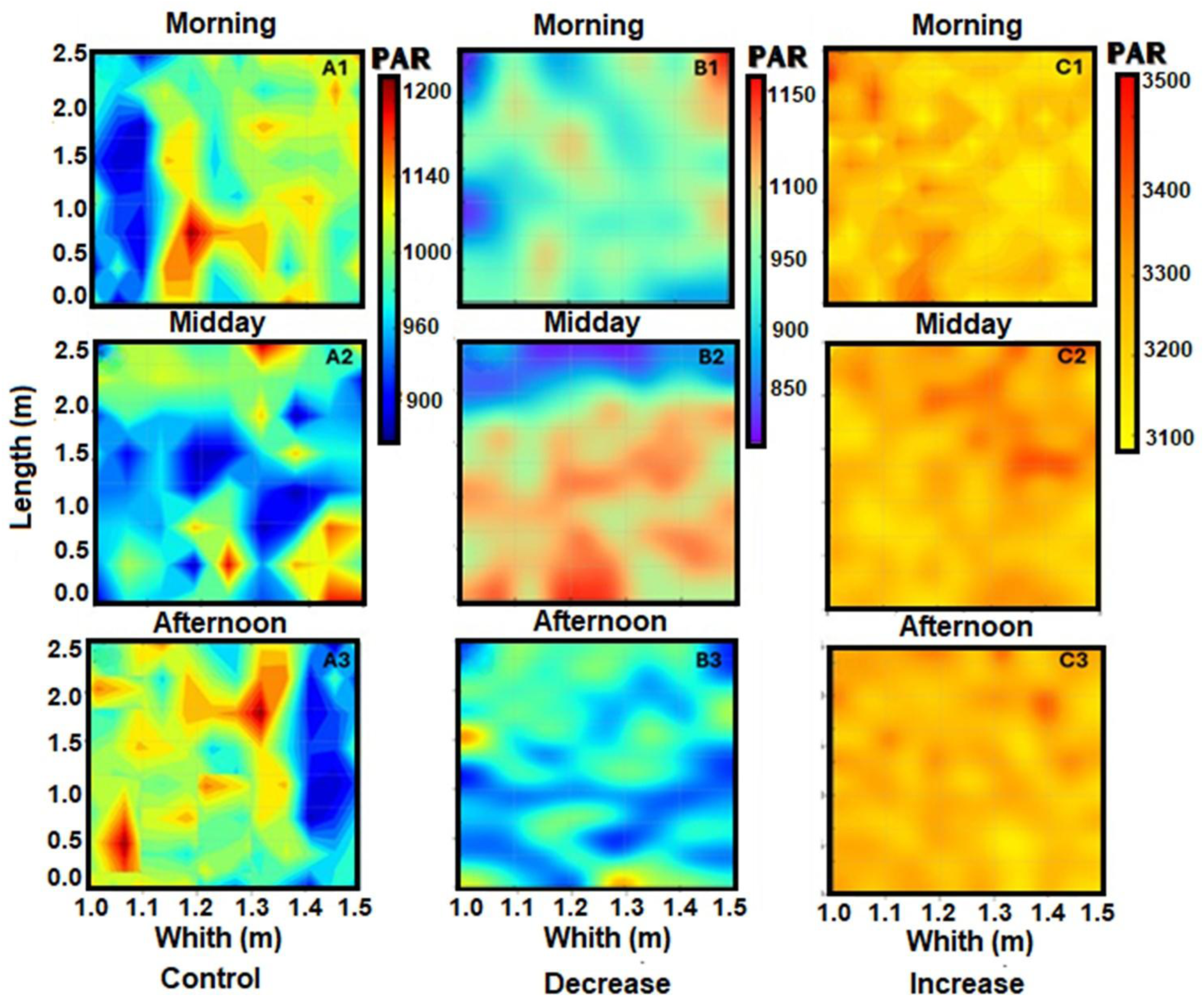
Spatial distribution of photosynthetically active radiation (PAR) measured at three times of the day (morning, midday, and afternoon). Panels A1–A3 show control PAR values, panels B1–B3 represent decreased PAR conditions, and panels C1–C3 display increased PAR conditions across the experimental plot. The color gradients indicate the intensity of PAR (red: high, blue: low).

### 2.4 Crop measurements

In both seasons, crop phenology was recorded twice a week following the BBCH phenological scale for rapeseed (Meier, 2001). The phenological stages included seedling emergence (BBCH 00-09), the beginning of flowering (BBCH 61), the end of flowering (BBCH 69), grain filling (BBCH 71), physiological maturity (BBCH 87), and harvest (BBCH 89). The crop cycle and developmental phases were expressed in thermal time units, calculated by summing daily mean temperatures using a base temperature of 0 °C (Kirkegaard et al., 2012).

Physiological maturity (BBCH 89) was determined when grains within siliques reached a dark color and hard texture in both seasons. At this stage, plant samples were collected from one linear meter of the central rows of each experimental plot. Biomass samples were separated into main stem, branches, siliques, and grains following the methodology described by Verdejo & Calderini, (2020). These samples were oven-dried at 65°C for 48 hours, and the individual weight of both siliques and grains was measured using an analytical balance (Mettler, Toledo XP205DR, Greifensee, Switzerland).

GY, harvest index (HI), GN, silique number, weight per silique, grains per silique, and TGW were measured or calculated. Aboveground biomass, GY, and silique weight were determined using a precision balance (Radwag WTC 2000, Radom, Poland), while grain number was obtained using a grain counter (Pfeuffer GmbH, Kitzingen, Germany). Harvest index was calculated as the ratio between GY and aboveground biomass. Silique weight was calculated by dividing the total weight of siliques by their number, while grain number per silique was determined as the ratio of the total grain number to the total silique number in the sample. The weight of thousand grains (TGW) was estimated by dividing GY by GN.

Oil concentration of grains was measured using Near-Infrared Reflectometry (NIR) (Foss Infratec 1241, Hilleroed, Denmark), and grain nitrogen concentration by the Kjeldahl method (Kirk, 1950). Protein concentration was determined on a dry matter basis using a conversion factor of 5.8, as recommended by Merrill and Watt (1973).

In season 2, five representative plants were sampled from each experimental plot to assess biomass accumulation and its distribution in vegetative and reproductive structures, at five phenological stages: BBCH 61 (beginning of flowering), BBCH 69 (full flowering), BBCH 71 (grain filling), BBCH 87 (physiological maturity), and BBCH 89 (harvest maturity). The collected samples were separated into main branches, secondary branches, siliques, and grains before being oven-dried at 65°C for 48 hours to ensure uniform processing, as described by Labra et al. (2017) and Verdejo & Calderini, (2020). Biomass accumulation dynamics, branch number, and dry matter were recorded as functions of thermal time after flowering (°C Day). Additionally, the weight of stems, branches, inflorescences and silique per plant were measured to evaluate structural and reproductive contributions.

### 2.5 Measurement of flower and individual grain dynamics

To account for flower and grain weight dynamics, these organs were measured from the beginning of flowering (BBCH 61) until harvest maturity (BBCH 89) in both experimental seasons. Flower measurements were taken on the main raceme and secondary branches, with samples collected from the beginning to the end of flowering (BBCH 69). The collected flowers were oven-dried at 65°C for 48 hours and weighed in the same analytical balance already mentioned to determine their dry weight.Grain measurements were carried out at specific positions on selected branches (branches 1 and 4). At each plot, 18 siliques were sampled, i.e. three siliques per position (basal, middle, and apical) from each of the two branches. Measurements were taken twice a week on marked plants, ensuring that no vegetative structures were removed during the sampling process. Siliques were also collected from the beginning of flowering to harvest maturity to evaluate the impact of the S-S treatments on final individual grain weight, grain growth rate, and duration. Grains from siliques were weighted immediately after collected to measure fresh weight using an analytical balance (Mettler, Toledo XP205DR, Greifensee, Switzerland). After that, grains and siliques without grains were oven-dried at 65°C for 48 hours, and the dry weight of grains and siliques were recorded in the same analytical balance.

The time course of grain weight was assessed at each branch position during grain filling by fitting a tri-linear broken-stick function, as described by Verdejo & Calderini, (2020).

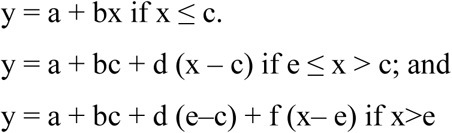

where grain weight (y) was estimated as a function of time (x) after anthesis (°C Day⁻¹) through three phases: the initial low dry matter accumulation phase (lag phase) with a grain filling rate (b), the linear accumulation phase with a linear grain filling rate (d), and a final phase with a reduced rate. The transition phase, i.e. break-points, were defined at times c and e, optimizing the fit to ensure continuity at the intersection points between segments. Comparisons of individual grain weight relative (%) to the control were calculated for both the reduced and increased S–S treatments, at branches 1 and 4, allowing a standardized evaluation across treatments and branches. Additionally, grain weight was evaluated in relative terms, calculated as the grain weight measured at each grain filling time relative to the final weight at physiological maturity (PM), thereby expressing the filling proportion in each treatment and experiment. The crop cycle and developmental stages were expressed in thermal time units (TT), calculated by summing daily mean temperatures (Tmean) with a base of 0°C (Kirkegaard et al., 2012), allowing assessment of biomass accumulation dynamics under the S–S conditions.

### 2.6 Statistical Analysis

The effects of the S-S treatments on biomass accumulation dynamics and crop properties were evaluated through detailed statistical analyses. Daily average differences in PAR and temperature among treatments were compared using an extra-sum-of-squares F-test in Stat graphics Centurion 18. The PAR distribution within the canopy for each plot during the treatment period was graphically represented using R software, specifically the ggplot2 and viridis packages, while temperature distribution was analyzed using GraphPad Prism 8.

Crop traits, including yield, yield components and grain quality (oil and protein concentrations in grains), were analyzed using a mixed-model analysis of variance (ANOVA) in Stat graphics Centurion 18. In this model, “experiment” was treated as a random factor, while “treatment” (S-S manipulation) and “branch position” were considered fixed factors. Significant differences between treatments were identified at a 5% probability level using the least squares mean differences test.

Linear regression analyses were performed with GraphPad Prism 8 to assess the data fit and to compare slopes and break-points of the linear relationships between grain weight, grain water content, and thermal time during the grain-filling period. The evaluation of grain weight was conducted both in absolute (mg) and relative (%) terms, consolidating the analysis for consistency across treatments A tri-linear segmented regression model was applied to fit the time-course of grain weight, allowing the identification of phase transitions (break-points) in biomass accumulation dynamics within grains, following the methodology described by Calderini and Reynolds (2000). Additionally, a third order centered polynomial model was used to adjust grain water content, as proposed by Menéndez et al. (2019), to prevent computational issues such as overflows (Motulsky, 2016). To ensure the validity of statistical inferences, comparisons of slopes and breakpoints were conducted across treatments and seasons. The assumptions of normality and homogeneity of residuals were verified using standard diagnostic tools, following the guidelines of Kutner et al. (2004).

## 3. Results

### 3.1. Crop phenology across experiments and environmental conditions

The BBCH 00–60 phase duration accounted for 46 days in the control treatment averaged across both seasons, accumulating 468°Cd (Fig. 2). Flowering and linear grain filling growth started at 532°Cd and 823°Cd, respectively, without differences (P > 0.01) among the S-S ratio treatments.

Mean temperature in control plots increased from 10.2°C (BBCH 00–59) to 14.1°C (BBCH 60–69) and 15.2°C (BBCH 70–89) along the crop cycle (Table 1). The reduced S-S ratio treatment lowered temperature by 1.5°C respect the control treatment, while the increased S-S ratio raised it by 1.3°C (Table 1). Similarly, midday silique temperature was reduced by up to 1.5°C under the reduced S-S ratio, while temperature of these organs was augmented by 1.3–1.5°C in the increased S-S ratio treatment (Fig. 3).

**Table 1.**
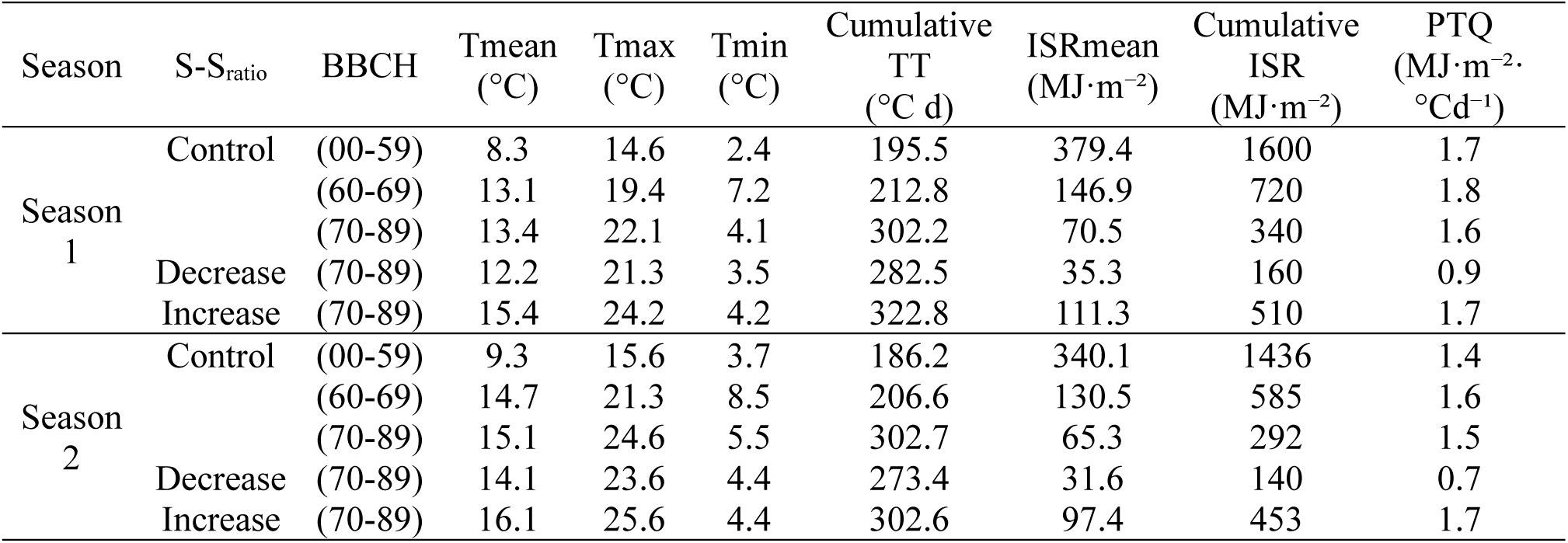
Mean, maximum and minimum temperatures (Tmean, Tmax and Tmin), cumulative thermal time (Cumulative TT), mean and cumulative incident solar radiation (ISRmean and Cumulative ISR), photothermal quotient (PTQ) during the phenological stages BBCH 00-59 (from seedling emergence to stem elongation), BBCH 60-69 (flowering), and BBCH 70-89 (silique development to ripening). The source-sink (S-S) ratio treatments (control, decrease and increase) were applied from the end of BBCH 60-69 (flowering) and continued through BBCH 70-89 (silique development to ripening) in Seasons 1 and 2.

However, these differences had little effect on the crop developmental phases, as physiological maturity was reached at ∼1050 °Cd in the control plots and at similar values (∼1000–1080 °Cd) in both source– sink (S–S) manipulation treatments (Fig. 2). Consistently, the thermal time duration of grain filling (BBCH 70–89) was 302 °Cd in the control and averaged ∼280 °Cd and ∼302 °Cd in both S–S treatments (Table 1), confirming that no significant (P > 0.05) differences were detected among the sources of variation.

Incident solar radiation in the control was 1,518 MJ m⁻² during the BBCH 00–59 phase averaged across both seasons, 653 MJ m⁻² from BBCH 60 to 69, and 316 MJ m⁻² between BBCH 70 and 89 along the crop cycle. The reduced S-S ratio decreased incident radiation by 50% below the control, while the increased S-S ratio improved it by 50% over the control, which was confirmed by radiation maps (Fig. 4). Despite variations in temperature and radiation, the photothermal quotient (PTQ) remained stable across treatments during BBCH 70–89, averaging 1.5 MJ·m⁻²·°Cd⁻¹ in the control. PTQ was reduced to 0.8 MJ·m⁻²·°Cd⁻¹ in the decreased S-S ratio treatment and augmented to 1.9 MJ·m⁻²·°Cd⁻¹ in the increased S-S ratio treatment, respect to the control (Table 1).

### 3.2. Grain yield, above ground biomass and harvest index

GY and biomass of the control treatment showed differences between seasons with the higher (P < 0.01) value in Season 1. In Season 2, both crop traits decreased by 32 and 28%, relative to the first season, respectively, due to lower cumulative radiation at flowering and higher temperature during grain filling (Table 1). These conditions were due to the later sowing date in Season 2, affecting biomass accumulation. In contrast, harvest index (HI) was similar between seasons (Table 2).

**Table 2.** Grain yield, above-ground biomass, harvest index, plant height, main stems weight and branches stem weight of source-sink (S-S) ratio treatments in Seasons 1 and 2.

| Season | S-S<sub>ratio</sub> | Grain yield<br>(Mg ha<sup>-1</sup>) | Above-ground<br>Biomass<br>(Mg ha<sup>-1</sup>) | Harvest<br>index<br>(%) | Plant height<br>(cm) | Main stems<br>weight<br>(g/plt) | Branches<br>stem weight<br>(g /plt) |
| --- | --- | --- | --- | --- | --- | --- | --- |
| Season 1 | Control | 7.5 ± 0.6 b | 25.4 ± 2.2 b | 30.2 ± 0.8 a | 171.3 ± 2.1 a | 2.5 ± 0.8 b | 1.2 ± 0.8 b |
|  | Decrease | 7.5 ± 0.9 b | 27.1 ± 2.5 b | 28.1 ± 0.7 a | 175.2 ± 3.0 a | 2.0 ± 0.7 b | 1.3 ± 0.7 b |
|  | Increase | 11.9 ± 0.8 a | 40.9 ± 3.0 a | 29.4 ± 0.8 a | 178.2 ± 3.1 a | 3.4 ± 0.5 a | 2.2 ± 0.5 a |
| Season 2 | Control | 5.1 ± 0.1 b | 18.3 ± 0.8 b | 29.6 ± 0.4 a | 177.7 ± 3.1 a | 2.1 ± 0.4 b | 1.1 ± 0.2 b |
|  | Decrease | 4.9 ± 0.2 b | 19.0 ± 0.5 b | 26.3 ± 0.8 b | 174.8 ± 4.9 a | 1.8 ± 0.7 b | 1.8 ± 0.7 b |
|  | Increase | 7.5 ± 0.2 a | 25.8 ± 0.5 a | 29.1 ± 0.3 a | 172.9 ± 3.8 a | 3.1 ± 0.5 a | 2.1 ± 0.4 a |
| Season (S) |  | < **0.001** | < **0.001** | 0.033 | 0.932 | < **0.001** | < **0.001** |
| S-S<sub>ratio</sub> (SR) |  | < **0.001** | < **0.001** | < **0.001** | 0.953 | **0.003** | < **0.001** |
| S x SR |  | 0.785 | 0.117 | **0.025** | 0.255 | 0.391 | 0.141 |

Mean ± standard error. Different letters indicate significant differences ( $p < 0.05$ ) between Season (S) and among source-sink treatments (SR). Significant p-values ( $p < 0.05$ ) are presented in bold.

Manipulation of the S–S ratio produced contrasting effects. The reduced source treatment had no impact (P > 0.05) on either GY or above-ground biomass but significantly decreased (P < 0.05) HI by about three percentage points under the reduced S–S ratio in Season 2. In contrast, the increased S–S ratio improved GY by 53% and biomass by 51% on average across seasons, while HI remained stable (Table 2). Additionally, no interaction between growing season and S–S ratio was observed for either GY (P = 0.79) or biomass (P = 0.12), confirming the consistent treatment response across seasons (Table 2; Fig. 5). Plant height was not affected (P > 0.05) by the sources of variation. The structural biomass (main stem and branch stems) varied according to season and treatment (Table 2). On average, it represented 15–18% of the above-ground biomass and was lower (P < 0.001) in Season 2 than in Season 1. The S–S manipulation had contrasting effects on structural biomass. Under reduced S–S ratio, this biomass was not affected (P > 0.05), minimizing the impact of a lower assimilate availability on these organs (Table 2). Conversely, structural biomass was enhanced (P < 0.001) under increased S–S ratio, indicating that greater radiation availability favored vegetative growth. Compared to the control, main stem biomass increased by 36% and branch biomass by 75% in Season 1, and by 48% and 91% in Season 2, respectively (Table 2).

**Figure 5.**
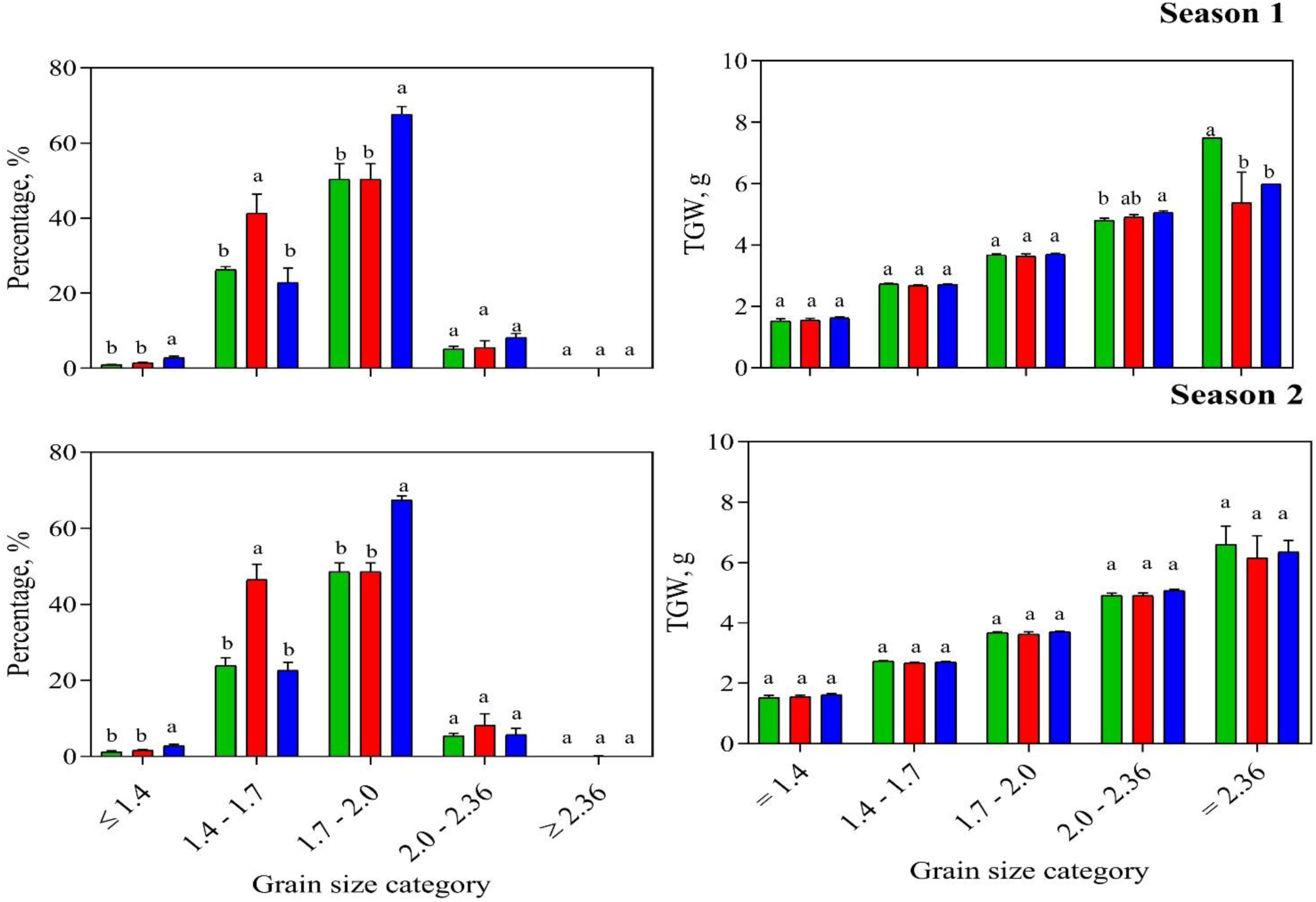
Frequency distribution (%) of individual grain weight across five grain size categories and its contribution to thousand grain weight (TGW) of control (green symbols and line), source decrease (blue symbols and line) and source increase (red symbols and line) treatments during seasons 1 and 2. Grain size was classified into five categories: <1.4, 1.4–1.7, 1.7–2.0,, 2.0–2.36 and >2.36 g.

### 3.3 Grain yield components

#### 3.3.1 Grain number

GN varied significantly between seasons and treatments (Table 3). Similar to GY and biomass, GN of the control treatment was higher in Season 1 and decreased by 27.6% in Season 2 (Table 3), likely due to the lower radiation during flowering that affected reproductive structures during Season 2 (Tables 1 and 3). Regarding S-S treatments, GN did not show (P > 0.05) difference under the decreased S–S ratio with the control treatment in Seasons 1 and 2 (Table 3). In contrast, the increased S–S ratio improved (P < 0.001) GN over the control by 56.5% and 40.5% in Seasons 1 and 2, respectively (Fig. A1). It is important to highlight that there was not (P = 0.180) interaction between season and S–S ratio, indicating a consistent treatment effect across years.

**Table 3.** Grain number (GN), thousand grain weight (TGW), siliques per plant, silique weight per plant and grains per silique of source-sink (S-S) ratio treatments in Seasons 1 and 2.

| Season | S-S<sub>ratio</sub> | GN<br>($10^3 \text{ m}^{-2}$) | TGW<br>(g) | Siliques per plant<br>(n) | Silique weight<br>per plant<br>(mg per plant) | Grains per<br>silique<br>(n) |
| --- | --- | --- | --- | --- | --- | --- |
| Season 1 | Control | 232 ± 19 b | 3.16 ± 0.03 b | 267 ± 32 b | 551 ± 48 b | 17 ± 2 b |
|  | Decrease | 210 ± 11 b | 3.46 ± 0.09 a | 259 ± 10 b | 523 ± 37 b | 16 ± 0.8 b |
|  | Increase | 363 ± 23 a | 3.26 ± 0.11 b | 383 ± 47 a | 834 ± 45 a | 24 ± 1.3 a |
| Season 2 | Control | 168 ± 7 b | 3.04 ± 0.11 b | 177 ± 32 b | 412 ± 35 b | 16 ± 2 b |
|  | Decrease | 150 ± 3 b | 3.53 ± 0.11 a | 159 ± 15 b | 394 ± 31 b | 15 ± 1 b |
|  | Increase | 236 ± 8 a | 3.18 ± 0.07 b | 253 ± 47 a | 724 ± 39 a | 21 ± 2 a |
| Season (S) |  | < **0.001** | 0.140 | < **0.001** | < **0.001** | **0.033** |
| S-S<sub>ratio</sub> (SR) |  | < **0.001** | < **0.001** | < **0.001** | < **0.001** | < **0.001** |
| S x SR |  | 0.180 | 0.576 | 0.117 | 0.083 | **0.025** |

Mean ± standard error. Different letters indicate significant differences ( $p < 0.05$ ) between Season (S) and among source-sink treatments (SR). Significant p-values ( $p < 0.05$ ) are presented in bold.

The impact of season and S-S treatments on GN was mainly explained by the number of siliques per plant, while grains per silique remained stable across treatments (Table 3). In the control plots, silique number decreased by 33.7% in Season 2. The main effect of S-S manipulations was found in the S-S ratio increase, where both siliques per plant and grains per silique were improved by 43% and 36% over the control, respectively, averaged across the seasons (Table 3)

The distribution of siliques along the plant profile followed a consistent pattern. In control plants, upper branches (B1–B3) supported more siliques than lower branches (B6–B7) (Fig. A2). The silique reduction in Season 2 was stronger in lower branches, with a 33.7% decrease compared to Season 1 (Fig. A2).

The S-S ratio treatments differently affected silique distribution. The reduction of the S-S ratio decreased the number of siliques per branch, with a stronger effect on lower branches, particularly in Season 2, i.e., 24% and 38% reduction compared to the control, respectively (Fig. A2). Conversely, when the S-S ratio was increased, silique production was improved, with the strongest effect on the upper branches (29% increase), followed by intermediate (18%) and lower branches (14%) in Season 1. In Season 2, these increases were 25%, 12%, and 10%, respectively (Fig. A2).

Another GN component, i.e., the number of grains per silique showed lower differences than the number of siliques per plant, ranging from 15 to 24 grains per silique across seasons and S-S treatments (Table 3). The higher values were recorded in Season 1 and under the S-S ratio increase. This treatment improved (P < 0.001) the trait by 41.2 and 31.2% over the control in Seasons 1 and 2, respectively (Table 3). These results highlight the number of siliques as the most sensitive component of GN, while the number of grains per silique was a more conservative trait, with significant but modest variation.

To have a better understanding of grain number generation, the weight of inflorescences was evaluated in the experiment (Fig. A3). Control plants allocated more inflorescence biomass in upper branches (B1– B3) across both seasons. In Season 2, the inflorescence weight was lighter, with a stronger reduction in the lower branches (−35% compared to Season 1). Similarly, the reduced S-S ratio led to a decline across branches, with stronger reduction in the lower ones (−24% in B1–B2 and −38% in B6–B7 relative to the control). On the contrary, the increased S-S ratio augmented the inflorescence biomass, showing increases of 29% in B1, 18% in B3, and 14% in B4 (Fig. A2).

#### 3.3.2 Grain weight

TGW was less sensitive than GN to both season and S-S ratio treatments. However, the decreased S–S ratio had a positive effect on this key trait as under shading TGW was increased (P > 0.05) over both the control and S-S increase by around 8% averaged across seasons, respectively (Table 3). This improvement was important to reach a similar GY between the S-S decrease and control treatments (Table 2).

In addition to TGW, grain size was assessed by sieving the harvested grains. The sieving process grouped grains into five categories (Fig. 5). In control plants, most grains were in the 1.7–2.0 mm and 2.0–2.36 mm classes. Under the reduced S–S ratio, TGW increased (P < 0.001) and grain size distribution shifted toward larger grains, with a higher proportion of grains above 2.0 mm at the expense of smaller categories. On the other hand, the increased S–S ratio reached similar distribution (P > 0.05) than the control (Fig. 5). In addition, TGW of the categories was similar across S-S treatments with the exception of >2.36 mm class of the control in Season 1.

When the impact of S–S ratio treatments on individual grain weight dynamics was assessed in basal, middle, and apical grains, all treatments showed a progressive filling pattern reaching physiological maturity between 870 and 910 °Cd (40–45 DAF) (Fig. 6). In control plots, grain filling proceeded at 0.074 g d⁻¹, with final weight stabilizing at ∼3.1 g. Under the reduced S–S ratio, basal and middle grains gained slightly more weight, increasing the grain-filling rate to 0.079 g d⁻¹ and delaying maturity to 910°Cd (45 DAF), which compensated the lower assimilate availability. In contrast, the increased S–S ratio enhanced the grain filling rate to 0.083 g d⁻¹, with maturity reached earlier at 870 °Cd (40 DAF), indicating faster assimilate accumulation. Under this treatment the highest final grain weight was achieve in basal grains, in line with the overall increases observed across positions (∼3.2–3.3 g). Detailed analyses by branch and grain position are provided in the Supplementary Information (Tables A2–A3, Fig. A5), confirming similar trends.

**Figure 6.**
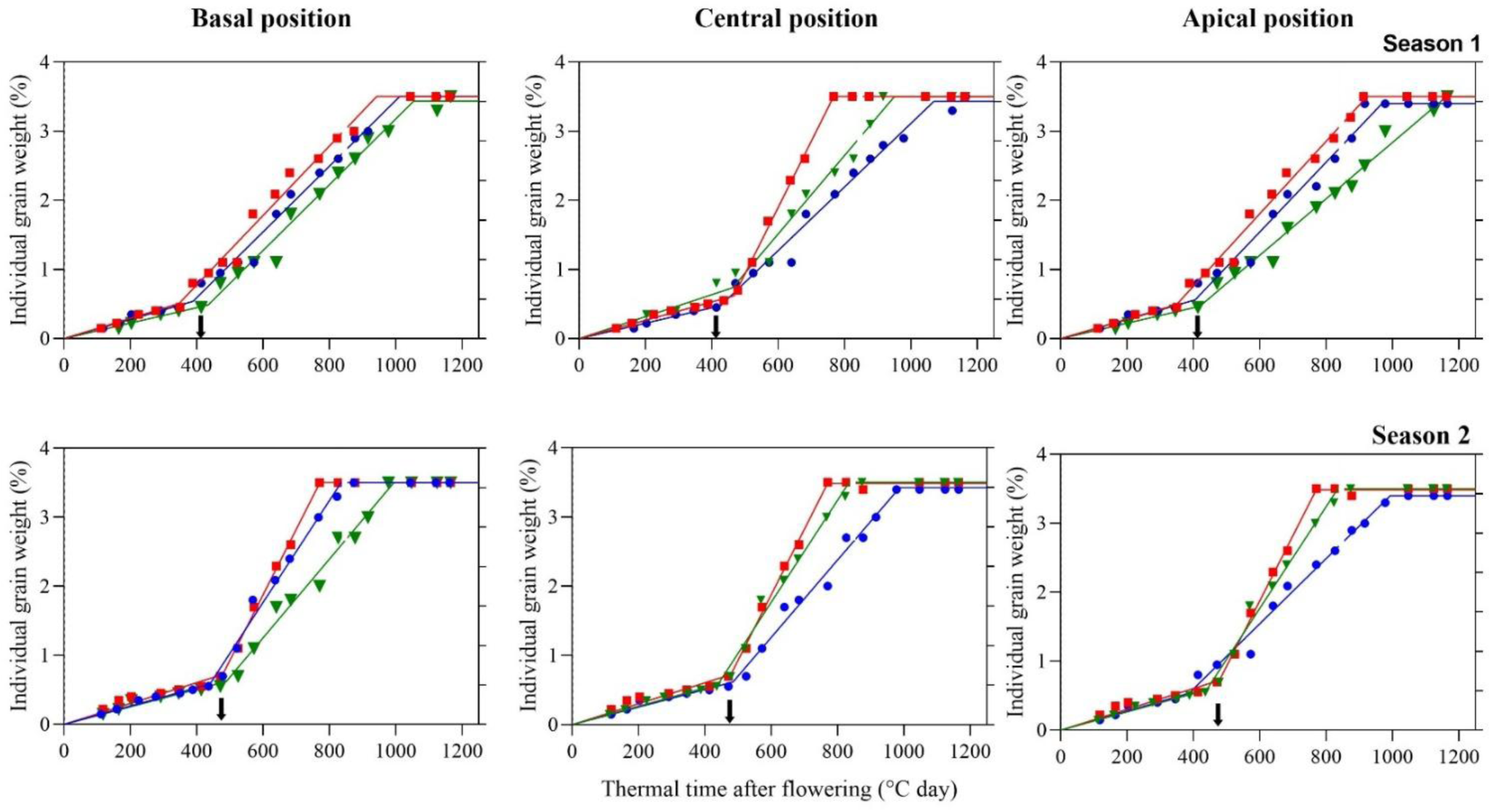
Individual grain weight (%) at basal, central, and apical positions during thermal time after flowering (°Cd) across two years. The phenological stages are divided into BBCH 60–69, corresponding to the critical period for grain number determination (Kirkegaard et al., 2018), and BBCH 70–89, corresponding to the grain-filling phase, during which the source-sink ratio treatments were evaluated in branch 1. Treatments are represented by colors: control (green), source increase (red), and source decrease (blue). In the first experiment, the grain-filling phase began at 412 °Cd, while in the second experiment, it began at 442 °Cd. Time points are indicated in cumulative degree days (°Cd). Arrows show the time when source-sink ratio (S-S_ratio_) treatments started.

### 3.4 Grain quality

As expected, grain oil concentration was as a quite conservative trait across seasons and S-S treatments, however effect (P < 0.001) by the sources of variation was found. In control plots, this quality trait averaged 50.2%, with a slightly higher value in Season 1 (50.8%) compared to Season 2 (49.6%) (Table 4). Under reduced S-S ratio, oil concentration decreased (P < 0.05) in both seasons, reaching an average of 48.6%, i.e., a reduction of 2.7 and 1.9 percentage points compared to the control in Seasons 1 and 2, respectively (Table 4). This means an effect, though mild, of the decreased source of assimilates on the oil synthesis. In contrast, the increase of the S-S ratio did not affect oil concentration in either season, although the increased source of assimilates improved GY by 55% (Tables 2 and 4).

**Table 4.** Grain oil protein concentrations under different source-sink (S-S) ratio treatments in Seasons 1 and 2.

| Season | S-S<sub>ratio</sub> | Grain oil concentration | Grain protein concentration |
| --- | --- | --- | --- |
|  |  | (%) | (%) |
| Season 1 | Control | 50.8 ± 0.4 a | 15.54 ± 0.40 b |
|  | Decrease | 48.1 ± 0.6 b | 17.92 ± 0.23 a |
|  | Increase | 50.9 ± 0.2 a | 16.35 ± 0.40 b |
| Season 2 | Control | 49.6 ± 0.3 a | 15.89 ± 0.58 a |
|  | Decrease | 47.7 ± 0.2 b | 16.24 ± 0.63 a |
|  | Increase | 48.8 ± 0.3 a | 15.71 ± 0.29 a |
| Season (S) |  | < **0.001** | 0.097 |
| S-S<sub>ratio</sub> (SR) |  | < **0.001** | **0.020** |
| S x SR |  | 0.092 | 0.125 |

Mean ± standard error. Different letters indicate significant differences ( $p < 0.05$ ) between Season (S) and among source-sink treatments (SR). Significant p-values ( $p < 0.05$ ) are presented in bold.

Another key quality trait of this crop, i.e. grain protein concentration, remained similar between seasons in control plants, averaging 15.7% across seasons (Table 4). When the S-S ratio was reduced, protein concentration increased (P < 0.05) by 2.4 percentage points only in Season 1, when this treatment had the highest impact on oil concentration. On the other hand, the increased S-S ratio had not affected grain protein concentration (Table 4). A trade-off between both quality traits was observed as a negative association was found when oil and protein concentrations were plotted (Fig. 7).

**Figure 7.**
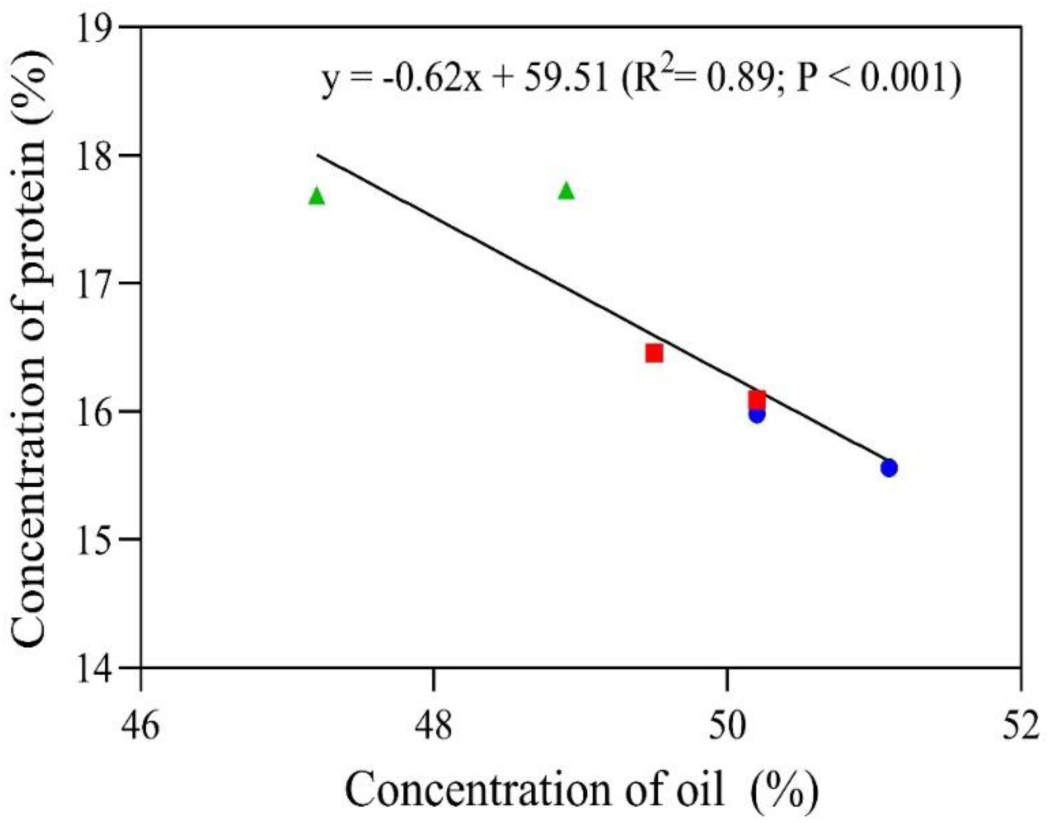
Relationship between grain oil and protein concentrations of control (green symbols and line), source decrease (blue symbols and line) and source increase (red symbols and line) treatments during Seasons 1 and 2.

## 4. Discussion

The aim of the present study was to contribute to answer the question if rapeseed is either source or sink limited during grain filling, regarding the controversial results reported in the literature. The manipulation of the S–S ratio in our study revealed a remarkable physiological resilience of rapeseed under a 50% reduction of incident solar radiation during the grain-filling phase (BBCH 71–89, 40–45 days after first flowering) in the high-yield environment of southern Chile. On the other hand, the rapeseed hybrid was highly sensitive to the increased S–S ratio by the supplemented solar radiation (50%), additionally oil and protein concentrations of grains were influenced by the S–S ratio manipulation, however, stable across seasons in the experiment.

### 4.1 GY under S-S ratio reduction

The resilience of GY to the decreased S–S ratio observed in the present experiment with the Click CL hybrid was also reported in other genotypes (Lumen and Solar CL) when the S-S ratio was reduced at different periods between flowering and physiological maturity in the environment of southern Chile (Labra et al., 2017; Verdejo & Calderini, 2020; Rivelli et al., 2024). This behavior reflects a compensatory capacity of the crop, whereby reductions in GN are partially or completely offset by increases in TGW, as has been reported under field conditions (Labra et al., 2017; Kirkegaard et al., 2018; Verdejo & Calderini, 2020; Rivelli et al., 2024). The stability of TGW observed here under source-reduction contrasts with studies focused on the grain-filling phase (Table A1) but it is consistent with the notion that source limitation is not a general condition for effective grain filling (Savin et al., under review).

Differences between experimental environments help to explain these contrasting results. Much of the evidence for strong decreases in TGW and oil concentration under source reduction comes from greenhouse or chamber studies, where shading of siliques accentuated environmental source limitations (Diepenbrock & Geisler, 1979; Fortescue & Turner, 2007). In contrast, field experiments have reported more heterogeneous responses. For instance, Pechan & Morgan (1985) found that basal defoliation at BBCH 71–75 led to variable reductions in GN and TGW, while Kirkegaard et al. (2012) showed that defoliation at BBCH 73–79 reduced seed number but not TGW, highlighting the sensitivity of seed set rather than grain growth to source limitation. Similarly, Mendham et al. (1981) and Gómez & Miralles (2010) emphasized that the magnitude of the response to source limitation is strongly conditioned by the environment, which explains why under the field conditions of southern Chile —long photoperiods, high solar radiation, and temperate springs— rapeseed showed stable yield despite a 50% source reduction (Rivelli et al., 2024). In addition, the proportion between S-S ratio reduction and grain weight (GW) change must also be considered (Borras et al., 2004; Savin et al., under review).

In our experiment, compensations were evident at the structural level. Biomass was preferentially redistributed toward basal branches (B1–B3), which maintained the number of fertile siliques and stable grain weight. This hierarchical buffering mechanism within the branched architecture of rapeseed prioritized grain filling in early reproductive structures and has been previously described in terms of lower abortion rates of main stems and lower-order branches under stress (McGregor, 1981; Tommey & Evans, 1992; Diepenbrock & Geisler, 1979). Quantitatively, TGW increased in the second season by 16.1% under decreased S–S relative to the control, offsetting a ∼14.5% lower GN in absolute values (P > 0.05). This agrees with Habekotté (1993), who quantified a compensatory relationship between GN and TGW in rapeseed.

At the physiological level, this stability can be explained by alternative carbon sources. Photosynthesis of siliques and stems may contribute up to 50% of the carbon required during seed filling (Müller & Diepenbrock, 2006; Fortescue & Turner, 2007), while remobilization of stem carbohydrates has also been identified as a key mechanism for maintaining grain weight under reduced assimilate supply (Rivelli et al., 2024). Furthermore, the stability of TGW suggests buffering mechanisms at the maternal–filial interface. Li et al. (2018) demonstrated that silique size and function, regulated by CYP78A9-mediated signaling, coordinates assimilate allocation to seeds, and Herrera & Calderini (2020) showed that pericarp tissues in wheat contribute assimilates directly to developing grains.

Taken together, these results indicate that rapeseed possesses a strong compensatory capacity under source reduction, combining structural buffering at the branch level with physiological mechanisms of assimilate supply during grain filling. This explains the resilience of GY and TGW observed in the present experiment and supports the interpretation that source limitation is not a universal constraint under field conditions, but one that depends strongly on environmental context and genotype (Rondanini et al., 2017).

### 4.2 GY in response to S-S ratio increase

The increase of the S–S ratio through enhanced solar radiation led to a substantial improvement in GY, driven by a significant rise in grain number, while individual grain weight remained stable. Diepenbrock (2000) and Zhang & Flottmann. (2018) reported that increased assimilation availability during flowering and early grain filling improves the setting of flowers, siliques, and grains, whereas grain size remains relatively constant. In soybean, Morandi et al. (1988) showed that photoperiod extension after flowering doubled pod number per plant under long-day versus short-day conditions, resulting in a marked increase in grain number. Likewise, Kantolic & Slafer (2007) quantified this effect by photoperiod extension (14 h vs. 10 h) in soybean, reporting >75% increases in pod and grain number per area, counterbalanced by ∼20% reduction in grain weight. Similarly, Mathew et al. (2000) observed that post-flowering light enrichment increased both pods per plant and grains per pod in soybean (∼70% gains), while grain size changed little. These findings are consistent with the present study in rapeseed, where the increase of the S–S ratio by augmented incoming solar radiation on the crop by 55%, enhanced GY through both more siliques and grains per plant. This result provides helpful information for rapeseed breeding aimed at improving grain yield potential. Moreover, this could be extrapolated to other semi-indeterminate crops species such as grain legumes to expand GN under favorable environmental conditions without penalizing grain weight.

### 4.3 Grain quality traits under both S-S ratio reduction and increase

The impact of S–S treatments on oil crops during grain filling must consider quality traits such as grain oil and protein concentrations in addition to GY and yield components to have a complete and precise picture of the response of these crops to different S-S ratios. In the present study, grain quality traits were conditioned by the S–S ratio manipulation in both seasons, as in previous reports showing asymmetric responses to assimilate availability (Triboi & Triboi-Blondel, 2013). Furthermore, this study observed that grain oil and protein concentrations were modulated by the S–S ratio, reflecting the sensitivity of seed composition to assimilate availability. Under source reduction (–S), oil concentration decreased by 2.7 and 1.9 percentage points in each season, while protein concentration increased. This inverse relationship has been widely documented in rapeseed (Triboi & Triboi-Blondel, 2013; Zhang & Flottmann., 2018; Li et al., 2018) and reflects competition between carbon allocation to lipid biosynthesis and nitrogen remobilization to grains. By contrast, the increase of the S–S ratio (+S) has not affected (P > 0.05) either oil or protein concentrations.

The contrasting response of oil and protein grain concentration to - S may stem from a decoupling between carbohydrate and protein synthesis during grain filling and the persistence of nitrogen in vegetative tissues, which, under moderate carbon limitation, is efficiently mobilized to the grain (Triboi & Triboi-Blondel, 2013).

Structurally, the +S treatment promoted more efficient assimilation allocation throughout the canopy, improving yield in both basal and distal branches (Appendix A). Unlike the –S treatment, where morphological compensation was localized in lower branches (B1–B3), improvements under +S were more homogeneous. This suggests that under high source availability, the plant can activate sinks in upper canopy positions that would normally be constrained by lower irradiance. This pattern aligns with findings by Tommey & Evans (1992), McGregor (1981), and Diepenbrock & Geisler (1979), who identified a hierarchical gradient of resource allocation toward physiologically prioritized branches. In this case, the increased assimilate supply appears to have overridden this hierarchy, maximizing total reproductive potential.

TGW stability under +S conditions confirms the stability and high heritability of this trait contrary to grain number as it is widely recognized. This was consistent across seasons and supports the rapeseed. Interestingly, in our experiment under +S, grain number expansion is favored, whereas under restrictive conditions (–S), grain filling efficiency is promoted.

Across the seasons of the experiment, the stability of both grain quality traits taking into account the relative change in the S-S ratio (∼ 50% decrease and increase) provides helpful information for breeding programs trying to increase grain yield of rapeseed regarding time and cost of quality traits evaluations. The stability of both oil and protein concentrations as well as their negative association is in agreement with previous reports from southern Chile (Verdejo & Calderini, 2020; Labra et al., 2017). These findings also provide empirical evidence for simulation models to modelled source–sink dynamics in oilseed crops (Porker et al., 2025; Fortescue & Turner, 2007). Nonetheless, critical questions remain regarding the genetic regulation of sink strength and the specific role of non-foliar photosynthetic organs under stress conditions (Diepenbrock, 2000; Harker et al., 2016). Future research should address the molecular regulation of assimilate partitioning and assess genotypic responses to S–S manipulations as strategies to improve rapeseed resilience and productivity. Importantly, the change in seed composition occurs without reductions in grain weight or total yield, indicating that the effect reflects a metabolic adjustment in assimilating partitioning rather than a limitation of grain filling.

## 5. Conclusions

The manipulation of the source–sink (S–S) ratio during the grain-filling period (BBCH 71–89) in rapeseed revealed contrasting responses of grain number (GN) and thousand grain weight (TGW) to changes in assimilate availability. This divergence reflects the physiological plasticity of rapeseed, enabling it to adjust reproductive development under both limiting and favorable source conditions without compromising vegetative or reproductive integrity.

In the control treatment, a balanced S–S ratio ensured stable GN and TGW, reflecting efficient resource allocation across branches and silique positions. Under source reduction, TGW was increased, especially in basal grains of upper branches, suggesting assimilates redistribution and the activation of structural buffering mechanisms. This response was accompanied by a more extended flowering period and a longer grain-filling duration (+3.4%), allowing the crop to maintain grain size under reduced carbon availability.

In contrast, the increase of the source significantly enhanced GN without affecting TGW, indicating that rapeseed can expand its numerical sink without affecting its filling capacity regarding also that the S-S increase was maintained during grain filling. This condition was associated with a more synchronized flowering pattern and a little shortening of the grain-filling period (–1.1%). Grain composition slightly responded to S–S ratio manipulation: source reduction little decreased oil content but increased protein concentration, while source enhancement had no effect.

Overall, the results confirm that rapeseed yield during grain filling is primarily determined by sink capacity. The crop’s ability to maintain yield and grain quality through compensatory mechanisms and efficient resource redistribution underscores its resilience and adaptability to diverse production environments.

## 6. Author contribution

Sebastian Garcia: Conceptualization, data curation, formal analysis, investigation, methodology, visualization, writing - original draft. José F. Verdejo: Conceptualization, formal analysis, investigation, methodology, writing – review and editing. Daniel F. Calderini: Conceptualization, investigation, methodology, resources, supervision, writing – review and editing. All authors read and approved the final version of the manuscript.

## Acknowledgements

We thank Olaff Sass (NPZ-Lembke, Germany) and Erik von Baer (Semillas Baer, Chile) for kindly providing the seeds of the genotypes. We really appreciate technical support from the staff of the Austral Farming Experimental Station (EEAA) of the Universidad Austral de Chile. The present study was supported by Project FONDECYT 1170913 (Chilean Technical and Scientific Research Council, CONICYT/ANID) competitive grant. José F. Verdejo held a postgraduate scholarship from CONICYT (ANID) 2017-21171384.

---

**Figure A1.**
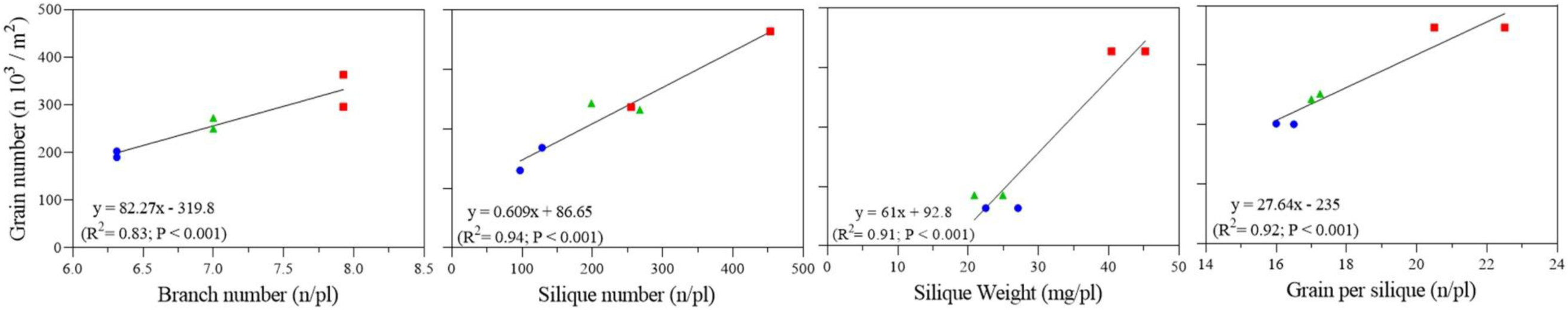
Relationships between grain number (n × 10³ m⁻²), branch number (n/plant), silique number (n/plant), silique weight (mg/plant), and grains per silique (n/plant) for two years. Treatments are represented by colors and symbols: control (green triangles), source increase (red squares), and source decrease (blue circles). Values represent the mean of four replicates per treatment. Data corresponds to season 1 and season 2.

**Figure A2.**
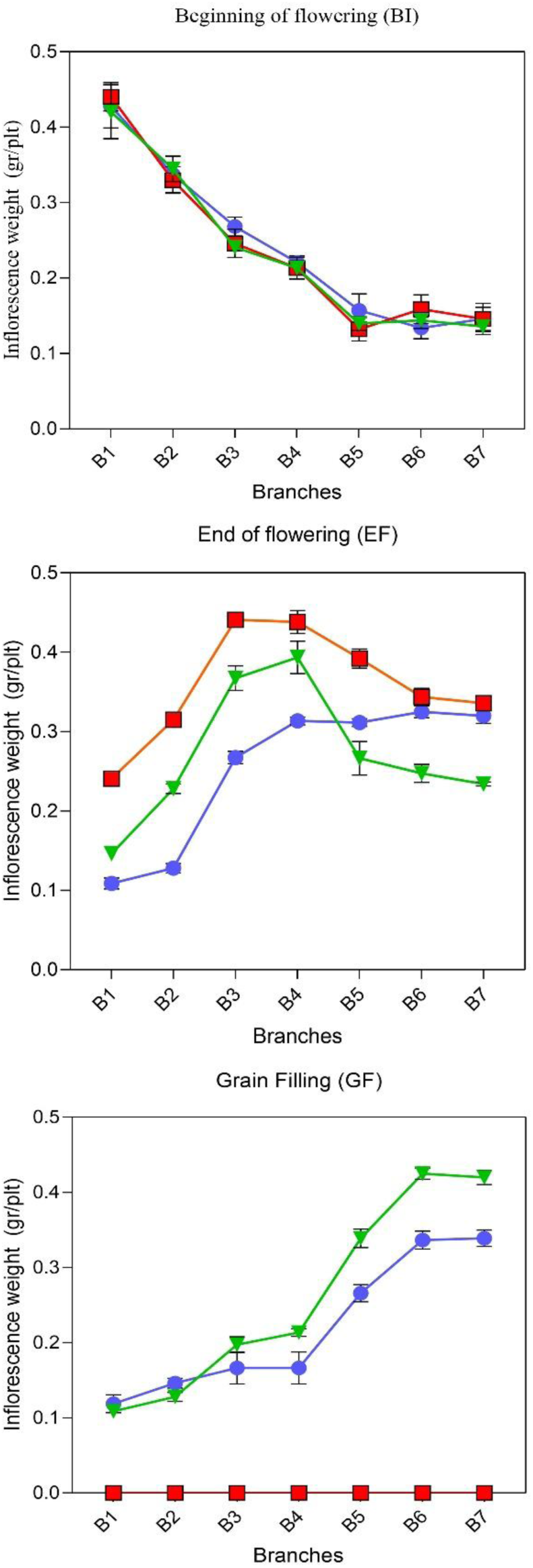
Dynamics of inflorescence weight (g/plant) across thermal time after flowering (°C d) in Experiment 2. Key phenological phases are indicated: beginning of flowering (400 °C d). end of flowering (600 °C d). and grain filling (1000 °C d). Branch-specific data are presented for branches B1 to B7. Treatments are represented by colors and symbols: control (green triangles), increase (red squares) and decrease (blue circles).

**Figure A3.**
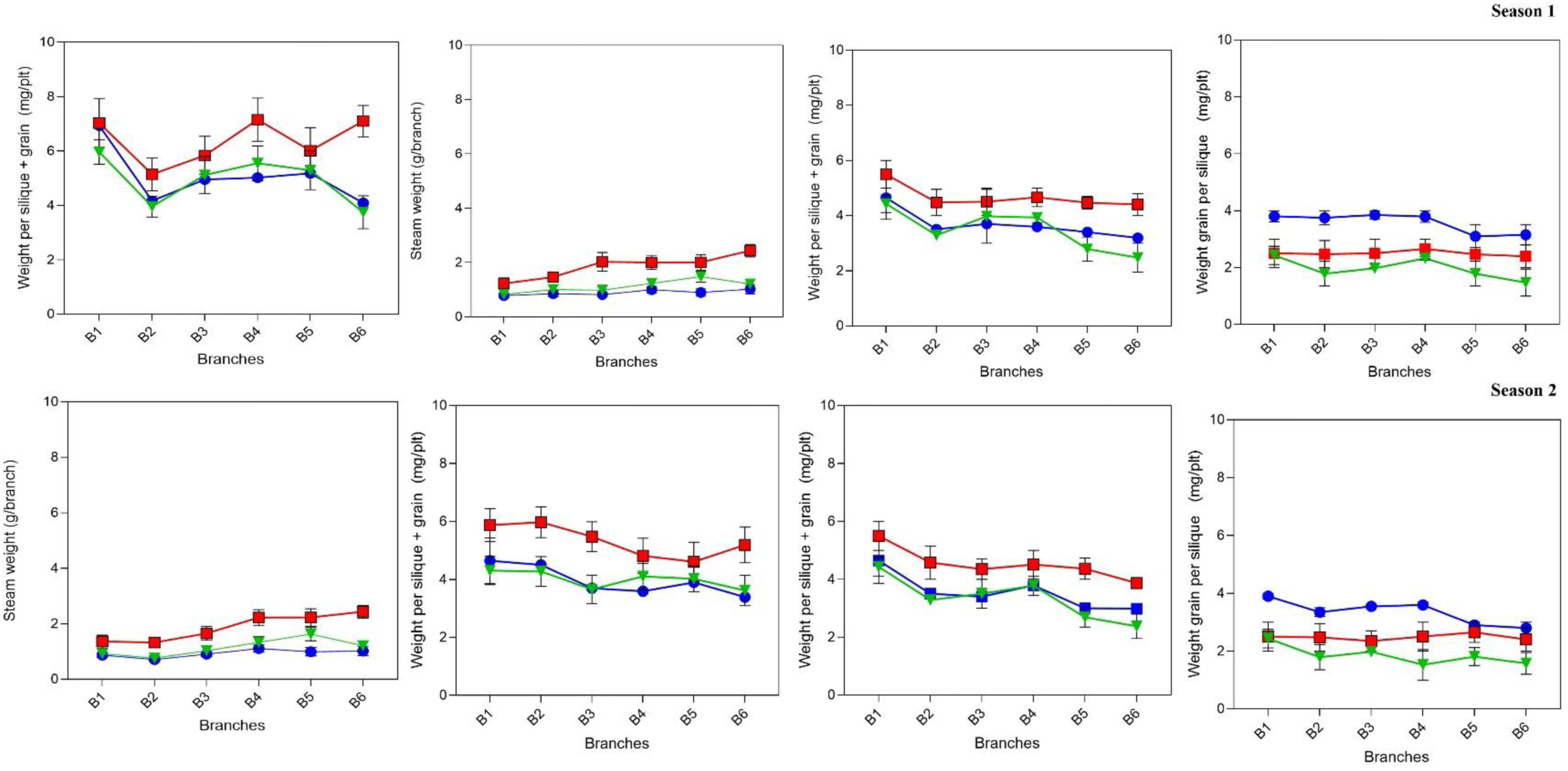
Stem weight (g/branch). weight per silique + grain (mg/pl). weight per silique (mg/pl). and weight of grains per silique (mg/pl) at harvest across branches B1 to B6. Treatments are represented by colors and symbols: control (green triangles). increase (red squares). and decrease (blue circles). Data corresponds to season 1 and season 2.

**Figure A4.**
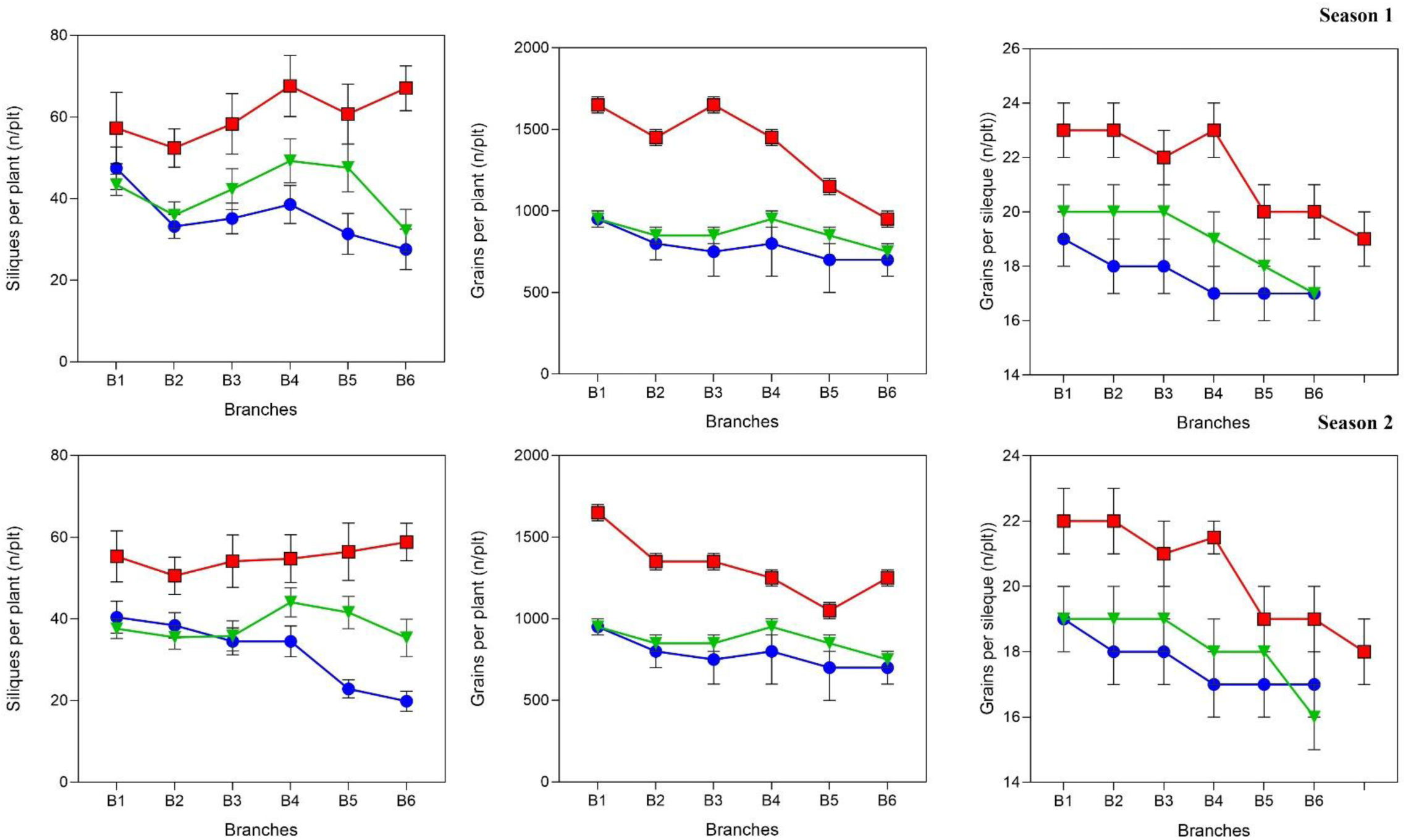
Siliques per plant (n/plant). grains per branch (n/plant). and grains per silique (n/plant) at harvest across branches B1 to B6. Treatments are represented by colors and symbols: control (green triangles). increase (red squares). and decrease (blue circles). Data corresponds to experiment 1 and experiment 2. Data corresponds to experiment 1 and experiment 2.

**Figure A5.**
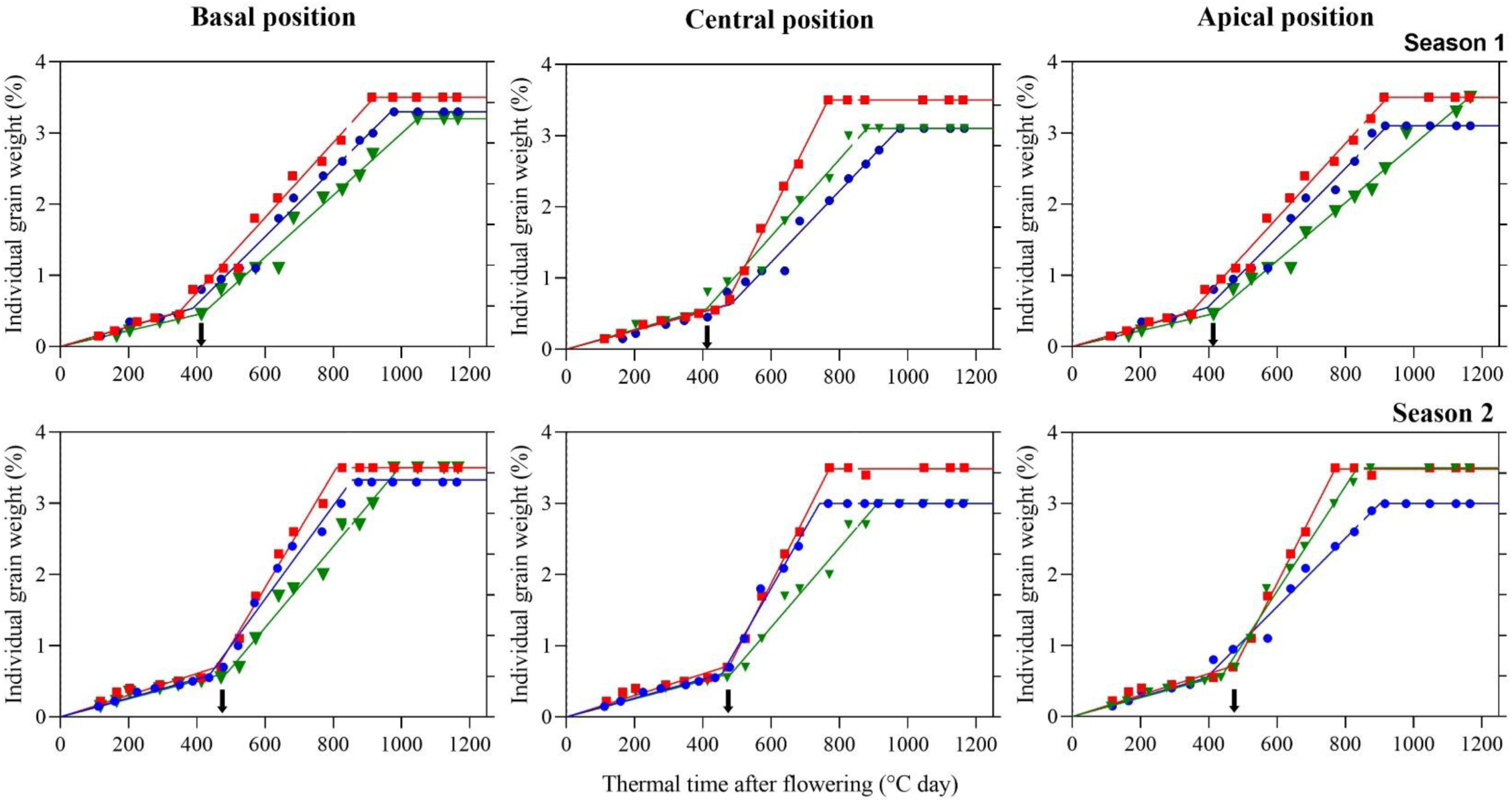
Individual grain weight (%) at basal, central, and apical positions during thermal time after flowering (°Cd) across two years. The phenological stages are divided into BBCH 60–69, corresponding to the critical period for grain number determination (Kirkegaard et al., 2018), and BBCH 70–89, corresponding to the grain-filling phase, during which the source-sink ratio treatments were evaluated in branch 4. Treatments are represented by colors: control (green), source increase (red), and source decrease (blue). In the first experiment, the grain-filling phase began at 412 °Cd, while in the second experiment, it began at 442 °Cd. Time points are indicated in cumulative degree days (°Cd). Arrows show the time when source-sink ratio (S-S_ratio_) treatments started.

**Table A1.**
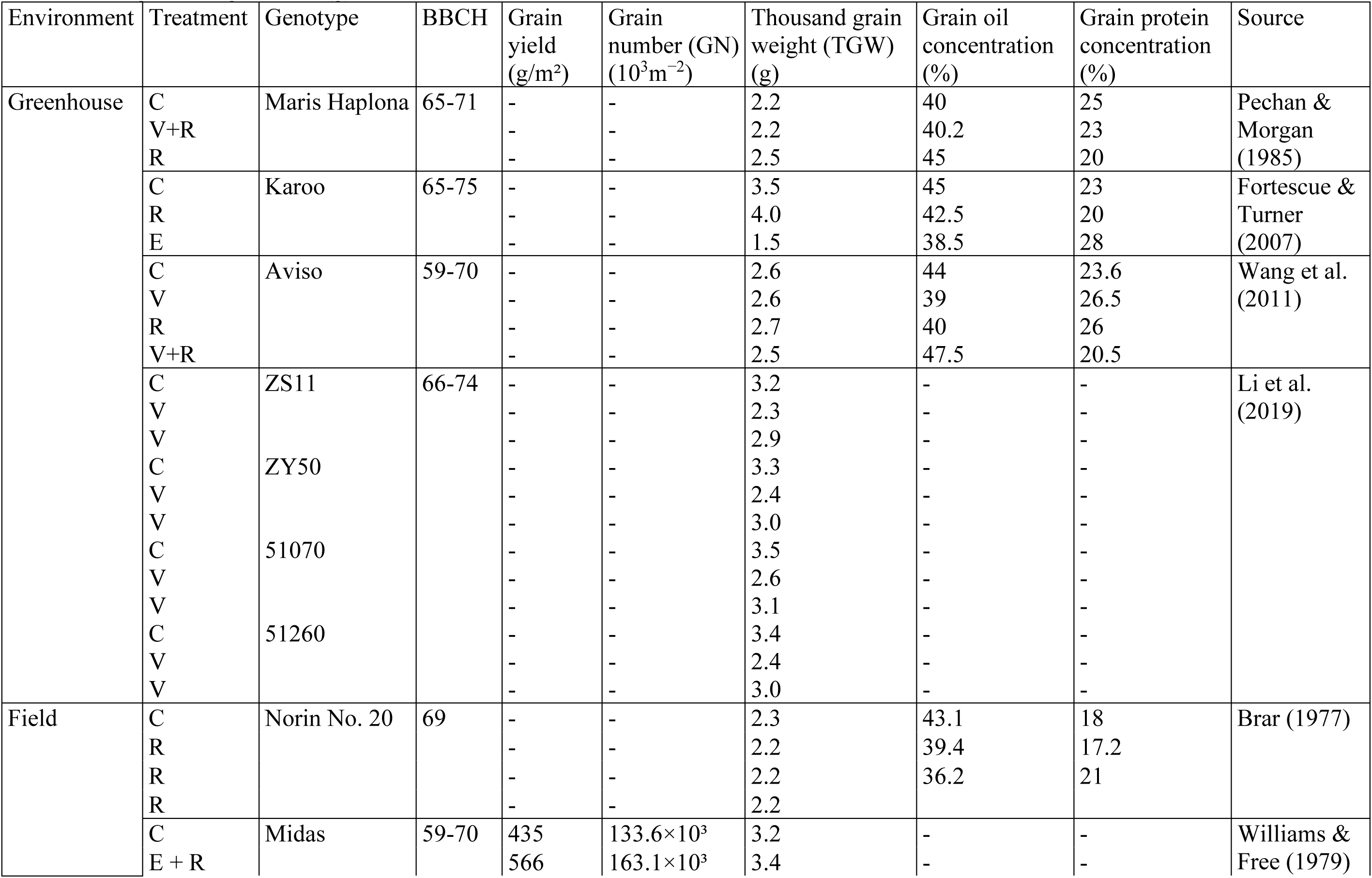
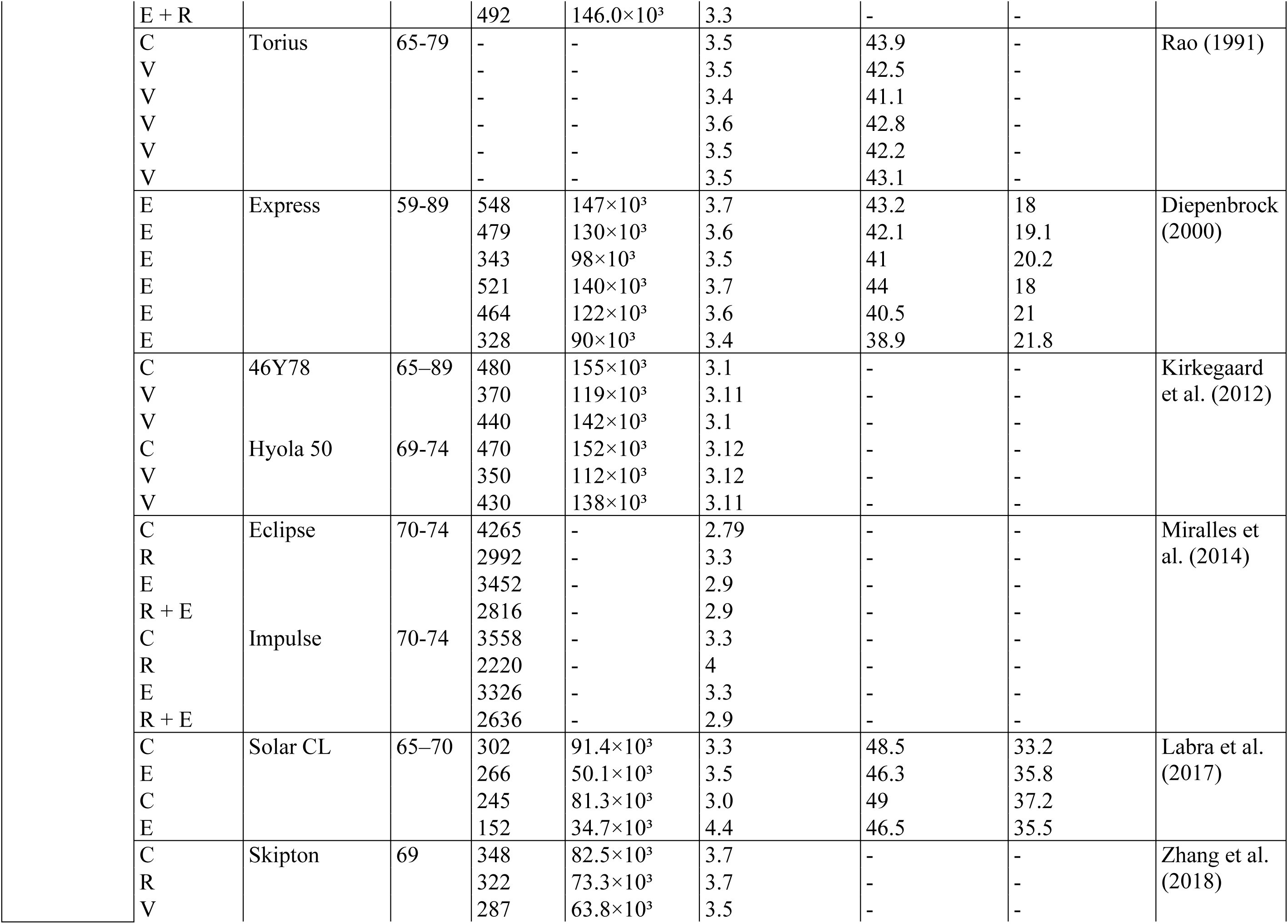
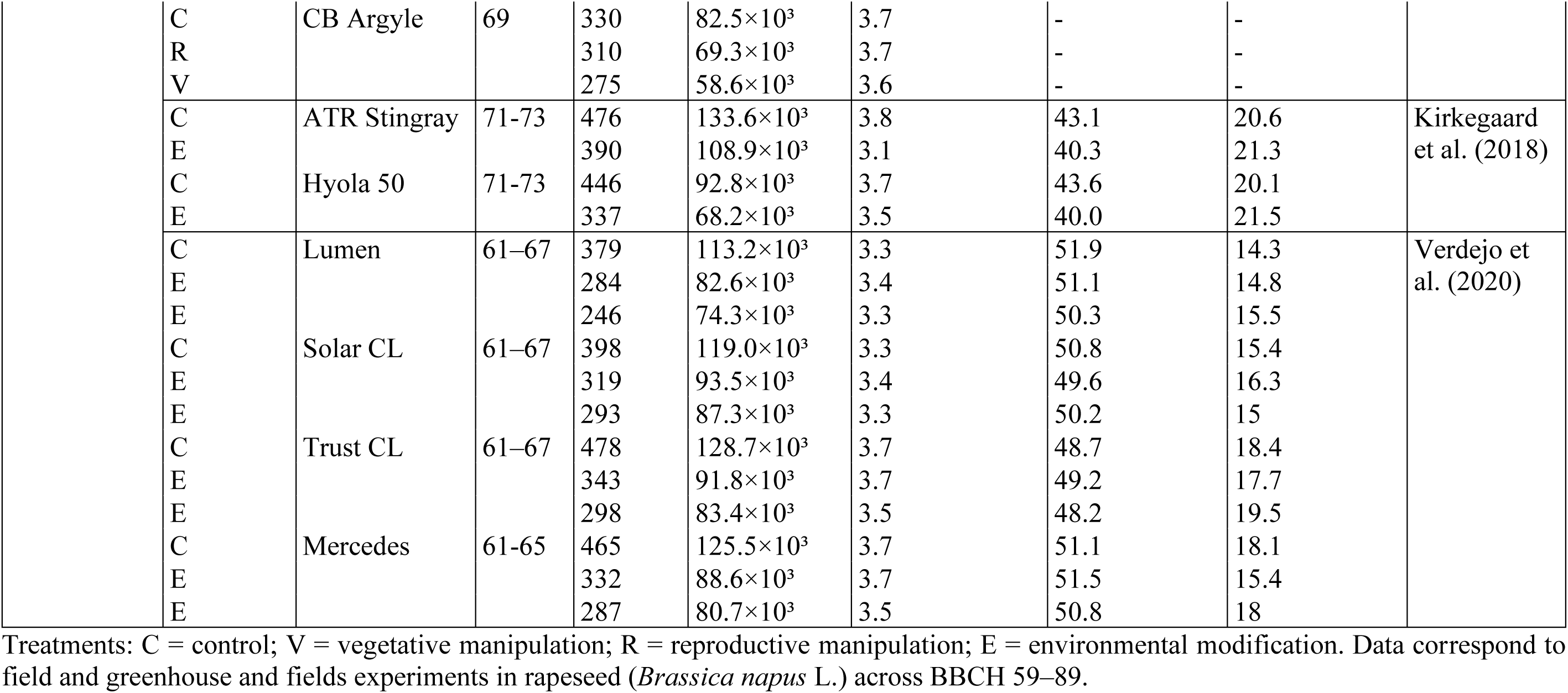
Comparative summary of studies assessing source–sink manipulations in rapeseed (*Brassica napus* L.) to evaluate compensation responses during grain filling. The table includes author, environment, treatment (source–sink manipulation), genotype, BBCH stage, grain yield (g/m²), grain yield per plant (g/plt), grain number per area (10³ grains/m²), grain number per plant (n/plt), grains per silique (n/silique), and thousand grain weight TGW (g)

**Table A2.**
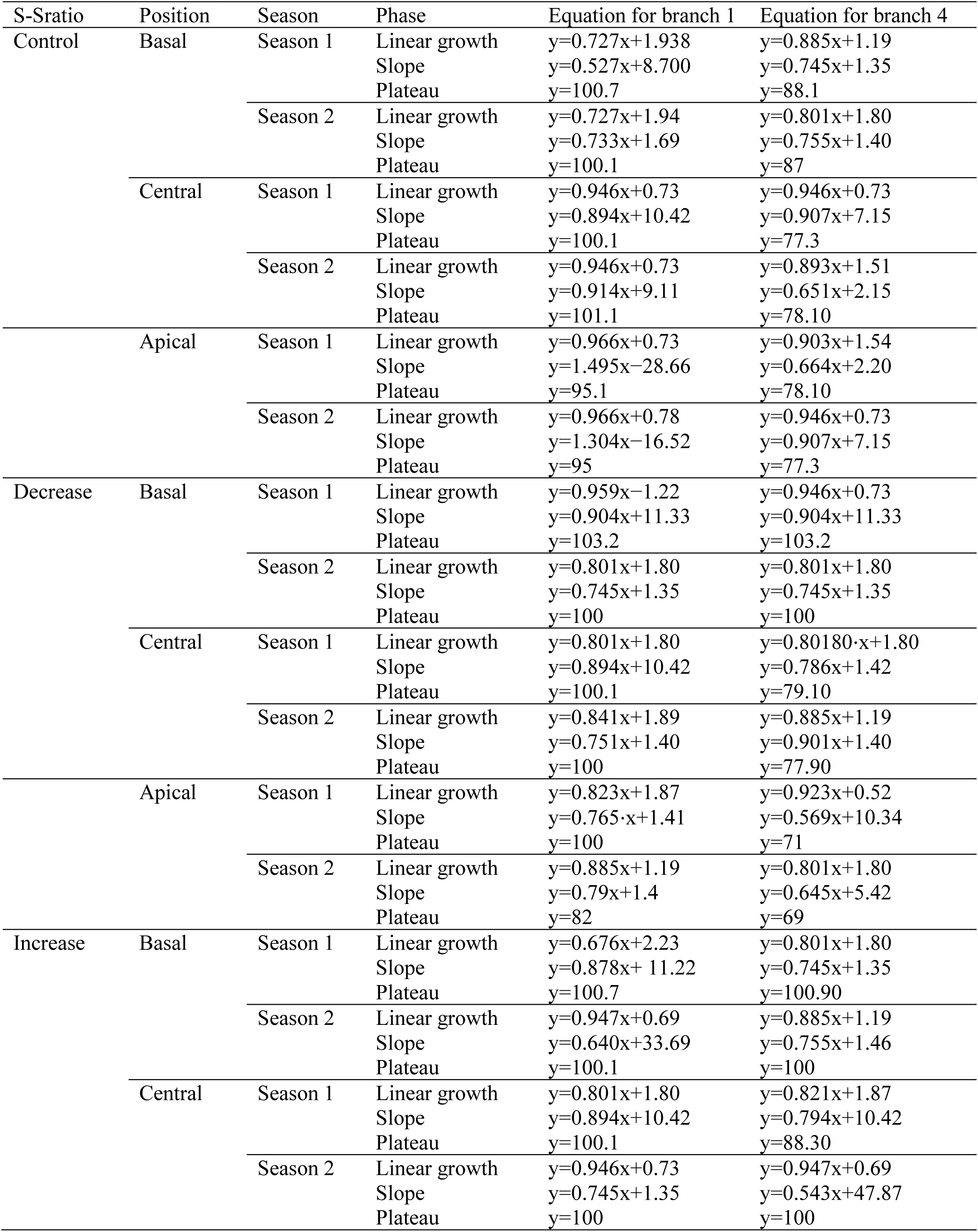
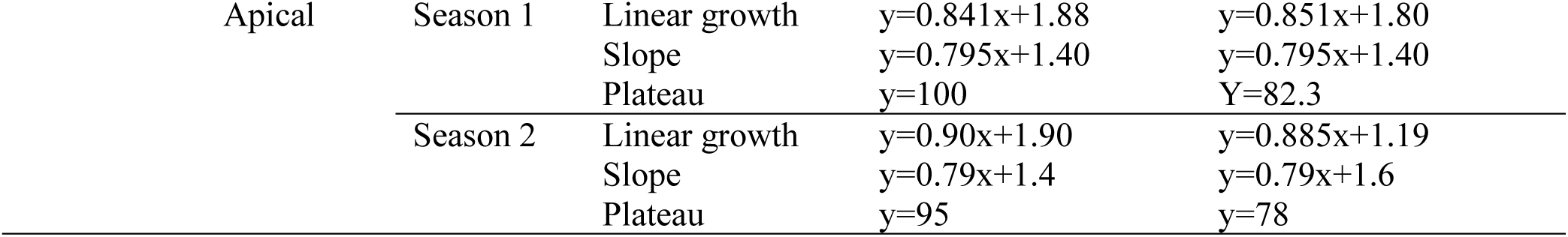
Equations of grain weight accumulation at basal. central. and apical positions across linear growth. slope and plateau phases in Experiments 1 and 2 for branch 1 and 4 from the end of flowering (BBCH 69) to harvest (BBCH 89).

## Notes

### Competing Interest Statement

The authors have declared no competing interest.

